# An allelic series reveals the genetic requirement for Adnp in cortical neurogenesis and learning behavior

**DOI:** 10.64898/2026.06.10.731333

**Authors:** Sarah Larrigan, Lina Dizon-Mapula, Iris Lasker, David J. Picketts, Pierre Mattar

## Abstract

Transcriptional regulators and chromatin remodellers are among the most important risk gene categories across the genetic landscape of neurodevelopmental disorders (NDDs). The zinc finger and homeodomain transcription factor *ADNP* is prominently associated with Helsmoortel-Van der Aa Syndrome (HVDAS), which is characterized by intellectual disability and autism spectrum disorder. Heterozygous frameshifting mutations account for the majority of HVDAS mutations, but it remains unclear how HVDAS mutations affect ADNP dosage, and how dosage in turn relates to neurodevelopmental and behavioral phenotypes. Here, we compared an allelic series of *Adnp* cKOs and germline heterozygotes. Using a conditional allele, we first deleted *Adnp* throughout the neural tube using *Nestin-Cre*. At E15.5, cKO brains exhibited altered upper-layer neuron production. However, *Adnp^Nestin^*cKOs exhibited perinatal lethality, precluding further behavioral characterization. Next, we compared germline heterozygotes (gHets) versus cKOs generated using the *Emx1-Cre* driver. We found that *Adnp^Del/+^*and *Adnp^L822fs6/+^* gHets exhibited cortical hypoplasia that was quantitatively identical, albeit less severe in comparison to *Adnp^Emx^*cKOs. In behavioral testing, *Adnp^Emx^* cKOs accordingly exhibited the strongest phenotypes, including hallmarks of elevated anxiety. However, both *Adnp^Emx^* cKOs and *Adnp^L822fs6/+^* gHets exhibited remarkably similar sex-specific deficits in learning during Morris Water Maze testing. Taken together, our work suggests that cortical growth and learning represent core phenotypes shared across *Adnp* mutant models irrespective of dosage. Moreover, since *Adnp* mutant phenotypes closely correspond with our prior findings in *Chd4* mutants, our results collectively suggest that Adnp regulates behavior via the ChAHP chromatin remodelling complex.

## Introduction

The mammalian neocortex plays a critical role in our ability to integrate sensorimotor information with higher order reasoning, which allows us to interpret our environment and surroundings with great sophistication. Accordingly, the neocortex has one of the most complex circuit structures found in the brain, with a host of different neuronal subtypes interconnected within a 6-layered columnar circuit. To generate different neuron subtypes, cortical progenitors must remodel gene expression dynamically in a stage-specific manner. Shifts in gene expression help direct the sequential production of deep-layer neurons, upper-layer neurons, and finally glial cells. Changes in gene expression and chromatin remodelling continue within differentiating daughter cells as they undergo migration, maturation, morphogenesis, and synaptogenesis.

In accordance with the requirement to dynamically remodel gene expression during lineage progression, chromatin remodellers have emerged as perhaps the most important neurodevelopmental disorder (NDD) risk gene category (De Rubeis et al., 2014; Iossifov et al., 2014; Valencia et al., 2023). However, in many cases, it remains unclear how genetic variants contribute towards behavior and NDD etiology. Moreover, since most chromatin remodellers are expressed ubiquitously in all cell types throughout the body, their gene regulatory functions are generally assumed to be generic rather than brain-specific.

The Activity-dependent neuroprotective protein (ADNP) is emblematic of these knowledge gaps. ADNP is a homeodomain and zinc finger transcription factor that is highly conserved in vertebrates (93% amino acid identity in human vs. mouse). Pathogenic *ADNP* variants are associated with Helsmoortel-Van der Aa Syndrome (HVDAS; also known as ADNP Syndrome) (Helsmoortel et al., 2014), which is associated with intellectual disability and speech delay. ADNP has also been identified as a high confidence autism risk gene (Satterstrom et al., 2020). Unbiased biochemical studies have revealed that ADNP forms a chromatin remodelling complex with CHD4 called ChAHP (<u>CH</u>D4, <u>A</u>DNP, <u>HP</u>1) that regulates the genome (Ostapcuk et al., 2018; Sharifi Tabar et al., 2022; Pintacuda et al., 2023). Perhaps accordingly, *CHD4* mutations are associated with Sifrim-Hitz-Weiss Syndrome (SIHIWES), which similarly features intellectual disability and autism spectrum disorder, as well as cortical growth phenotypes such as macrocephaly and microcephaly. SIHIWES and HVDAS thus share some phenotypic features, and they also share a very similar episignature (Sifrim et al., 2016; Weiss et al., 2016; Weiss et al., 2020; Karimi et al., 2025).

To identify the neurodevelopmental processes that depend upon *Adnp*, germline mouse mutants were previously generated. However, homozygotes were lethal at early embryonic stages (Pinhasov et al., 2003). To bypass this lethality, we previously created a conditional null allele by flanking exon 5 with loxp sites. Exon 5 encodes 94% of the ADNP protein including all of the DNA binding motifs (Clemot-Dupont et al., 2025). We then generated telencephalon-specific conditional knockouts (*Adnp^Emx^* cKOs) using the *Emx1-Cre* driver (Gorski et al., 2002). Notably, we reported that *Adnp^Emx^* cKOs and *Chd4^Emx^* cKOs were virtually indistinguishable, with both cKOs exhibiting cortical hypoplasia, reduced upper-layer neurons, and similar transcriptomic signatures. This phenotypic similarity suggests that Adnp regulates cortical neurogenesis through the ChAHP complex. However, a later requirement for Adnp and the ChAHP complex in particular circuits and behaviors has not yet been established. While conditional genetics approaches can potentially map phenotypes to specific brain regions and cell types, it additionally remains unclear to what degree *Adnp* cKOs correspond with germline heterozygous mutants that better mimic the genetics of HVDAS.

To begin to address these research gaps, we directly compared *Adnp^Emx^*cKOs with additional mutants, including cKOs generated using the *Nestin-Cre* driver (*Adnp^Nestin^*), as well as germline heterozygotes (gHets; *Adnp^Del/+^*) and mice harboring a C-terminal frameshfting CRISPR indel (*Adnp^L822fs6/+^*) (D’Incal et al., 2026). Using this allelic series, we report that *Adnp^Nestin^* cKOs exhibited cortical hypoplasia that closely resembles the phenotype previously observed in *Adnp^Emx^* cKOs. However, whereas *Adnp^Emx^*cKOs were viable and fertile, *Adnp^Nestin^* cKOs were perinatally lethal. *Adnp^Del/+^* and *Adnp^L822fs6/+^*gHets also exhibited gross reductions in brain size and weight. While brain hypoplasia is thus a common feature, its severity varied according to gene dosage. Next, we performed behavioral characterization. We found that *Adnp^Emx^*cKOs exhibited phenotypes exploratory behaviors, anxiety, fear and learning behaviors, with some differences appearing to be sex-specific. Most of these phenotypes were not observed in conditional heterozygotes (cHets), but equivalent deficits in learning were observed in *Adnp^Emx^* cKOs and *Adnp^L822fs6/+^* gHets during the Morris Water Maze test. Perhaps more interestingly, several of these behavioral phenotypes mirrored those of equivalent *Chd4* mouse models previously assessed using the same testing pipeline (Larrigan et al., 2023), suggesting that shared phenotypes may be driven by the ChAHP chromatin remodelling complex. Taken together, our work defines core neurodevelopmental phenotypes associated with *Adnp* mutations, including cortical growth, anxiety, and learning behaviors.

## Results

The ADNP protein includes 9 zinc finger motifs that span the N-terminus, followed by a well characterized nuclear localization sequence (NLS), and then a homeodomain that has been shown to be essential for genome occupancy (Yan et al., 2022). The HVDAS genetic landscape is predominated by frameshifting mutations distributed across the coding sequence, although hotspots are present around the NLS and homeodomain (Helsmoortel et al., 2014; Harutyunyan et al., 2026) (Fig. 1A). Since most of these mutations occur within the final exon of *Adnp* (which contains ∼94% of the coding sequence), nonsense-mediated decay mechanisms appear not to be triggered (Helsmoortel et al., 2014). However, investigations reported that truncated proteins were undetectable in cells harboring HVDAS mutations (Yan et al., 2022; D’Incal et al., 2024). These observations suggest that frameshifted proteins are oftentimes unstable, although it remains unclear whether this is systematically true for all nonsense mutations associated with HVDAS.

**Figure 1.**
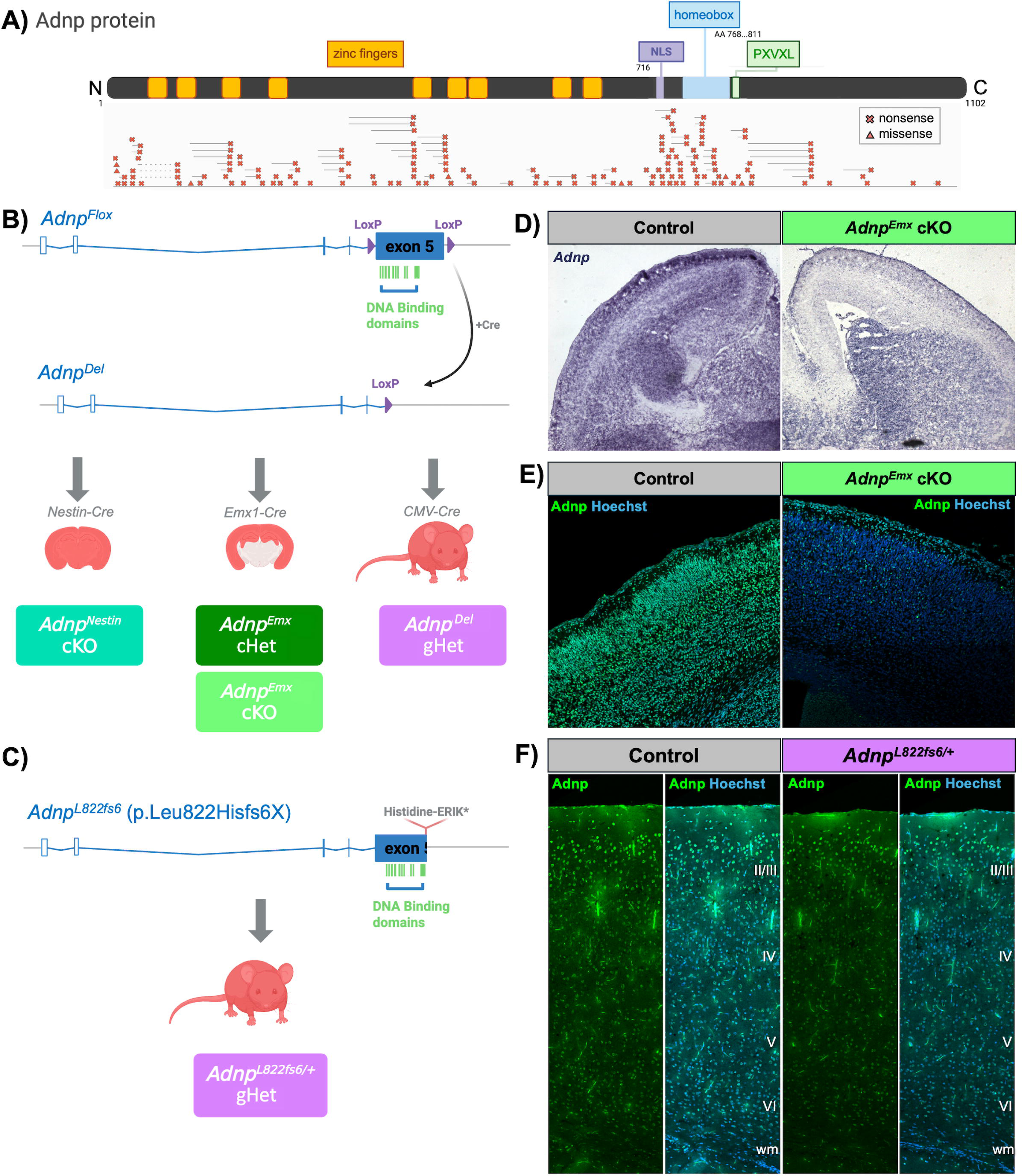
An allelic series of *Adnp* mouse models. (A) ADNP protein structure and pathogenic variants. N.b. scale approximate. (B) Genetic strategy for generating *Adnp^Nestin^*cKOs, *Adnp^Emx^* cKO/cHets, and *Adnp^Del/+^* gHets. (C) *Adnp^L822fs6/+^* gHets (D’Incal et al., 2026). (D) *Adnp* mRNA *in situ* hybridization, and (E) Adnp immunofluorescence on wild-type vs. *Adnp^Emx^*cKOs as indicated. (F) Adnp immunofluorescence on wild-type vs *Adnp^L822fs6/+^*gHets as indicated.

To gain insight into how differences in Adnp dosage affect neurodevelopment and behavior, we generated an allelic series of mouse models. Previously, we generated a strong conditional *Adnp* allele, in which exon 5 was flanked by loxp sites (Clemot-Dupont et al., 2025). To delete *Adnp* throughout the neural tube, we crossed the *Adnp^Flox^* allele with a *Nestin-Cre* driver that is active throughout the neural tube (Berube et al., 2005). We also examined cKOs in which *Adnp* was deleted specifically from the dorsal telencephalon-using the *Emx1-Cre* driver (Gorski et al., 2002). To generate germline heterozygotes, we additionally excised the ‘floxed’ cassette using the *CMV-Cre* driver (Schwenk et al., 1995) (Fig. 1B). Finally, we also examined germline heterozygous mice harboring a p.Leu822Hisfs*6 frameshifting CRISPR indel (*Adnp^L822fs6/+^*) designed to mimic C-terminally truncating *ADNP* variants associated with HVDAS (D’Incal et al., 2026) (Fig. 1C). Collectively, this approach generated an allelic series with different levels of predicted severity (in order of strongest to weakest): *Adnp^Nestin^*cKOs, *Adnp^Emx^* cKOs, *Adnp^L822fs6/+^* gHets, *Adnp^Del/+^* gHets, *Adnp^Emx^* cHets. As shown previously (Clemot-Dupont et al., 2025), *Adnp* mRNA and protein was efficiently deleted from the dorsal telencephalon (Fig. 1D, E), validating the conditional strategy. Using knockout-validated N-terminal antibodies, we did not observe a clear effect on Adnp protein expression in *Adnp^L822fs6/+^* gHets (Fig. 1F) in accordance with previous studies (D’Incal et al., 2026).

### Adnp is required in the CNS for viability and cortical neurogenesis

Prior work has shown that homozygous *ADNP* mutation arrests the induction of neural progenitors (Pinhasov et al., 2003; Ostapcuk et al., 2018; Sun et al., 2020b; Yan et al., 2022; Cho et al., 2023). By contrast, we previously showed that *Adnp^Emx^* cKOs were viable and fertile, although they exhibited cortical hypoplasia (Clemot-Dupont et al., 2025). Since *Emx1-Cre* expression is restricted to the dorsal telencephalon, we decided to first examine the consequences of deleting Adnp in a more widespread fashion using a CNS-wide *Nestin-Cre* driver allele that begins to be expressed at approximately embryonic day (E) 11.5 in the telencephalon (Berube et al., 2005). We examined juvenile mice post-weaning but failed to recover viable *Adnp^Nestin^* cKOs (p < 0.0001, chi-square test, n=57). Next, we harvested litters at E15.5. Embryonic *Adnp^Nestin^* cKOs were recovered in approximately Mendelian ratios, suggesting that cKOs likely die perinatally.

We next examined E15.5 *Adnp^Nestin^* cKO brains using immunohistochemistry. We found that Adnp protein was largely ablated throughout the cortical plate, with the exception of some neurons within the marginal zone and subplate, which are the earliest born neuronal subtypes (Fig. S1A). There also appeared to be residual perdurance of Adnp protein in the ventricular and subventricular zones of most *Adnp^Nestin^* cKOs, suggesting some variability in the efficacy of excision. Measuring sections, we found that the overall thickness (Fig. S1C) and cell density (Fig. S1D) of the cortex was not significantly altered in *Adnp^Nestin^* cKOs versus controls (consisting of cre-negative, as well as cre-positive *Adnp^+/+^*and *Adnp^Flox/+^* genotypes). Since upper-layer neurons were underproduced in *Adnp^Emx^*cKOs (Clemot-Dupont et al., 2025), we next examined neuronal subtypes. We noted a significant decrease in Pou3f2+ upper-layer neurons within the intermediate zone as well as in the superficial cortical plate (Fig. S1H). In contrast, we found little difference in early-born Bcl11b+ neurons between *Adnp^Nestin^*cKOs and controls (Fig. S1I). Taken together, these data indicate that that *Adnp^Nestin^* cKOs exhibit cortical neurogenesis defects that resembled the upper-layer defects previously reported in *Adnp^Emx^* cKOs (Clemot-Dupont et al., 2025). However, the observed lethality indicates that *Adnp^Nestin^*cKOs are intermediate between germline *Adnp* homozygous knockouts (lethal at ∼E8.5) (Pinhasov et al., 2003), and *Adnp^Emx^* cKOs (viable and fertile).

### Adnp^Del/+^ and Adnp^822/+^ germline heterozygotes exhibit growth defects

Next, we compared germline heterozygous models that better match HVDAS genetics. Examining gross anatomy, we found that both *Adnp^Del/+^*and *Adnp^L822fs6/+^* gHets exhibited decreased body weight and brain weight, which was more prominent in females (Fig. 2A-E). *Adnp^Del/+^* and *Adnp^L822fs6/+^* gHets also presented with decreased cortical area (Fig. 2F). However, this was not as drastic as previously observed in *Adnp^Emx^*cKOs (Clemot-Dupont et al., 2025) (Fig. 2C), and also correlated with reduced cerebellum area (Fig. 2G), suggesting a general growth defect. Next, we performed histological analyses on P28 cortices, focusing on the somatosensory cortex of *Adnp^L822fs6/+^* gHets (Fig. S2G, H). In accordance with previous work (D’Incal et al., 2026), *Adnp^L822fs6/+^*brains exhibited no significant change in cross-sectional thickness, cell density, number of Hoechst+ cells (Fig. 2H-J), nor in counts of upper layer (Bcl11b+) or lower layer (Pou3f2+) neurons (Fig. 2K, L). While cortical lamination was not quantitatively affected in *Adnp^L822fs6/+^* gHets, re-analysis of published RNA-seq data (D’Incal et al., 2026), revealed that key transcription factors associated with upper-layer identity were systematically downregulated in *Adnp^L822fs6/+^* gHets, while lower-layer markers were generally unaffected (Fig. S2). Along with prior work in *Tbr1* knockouts (Notwell et al., 2016), these observations collectively link Adnp to gene expression programs associated with upper-layer neuron specification.

**Figure 2.**
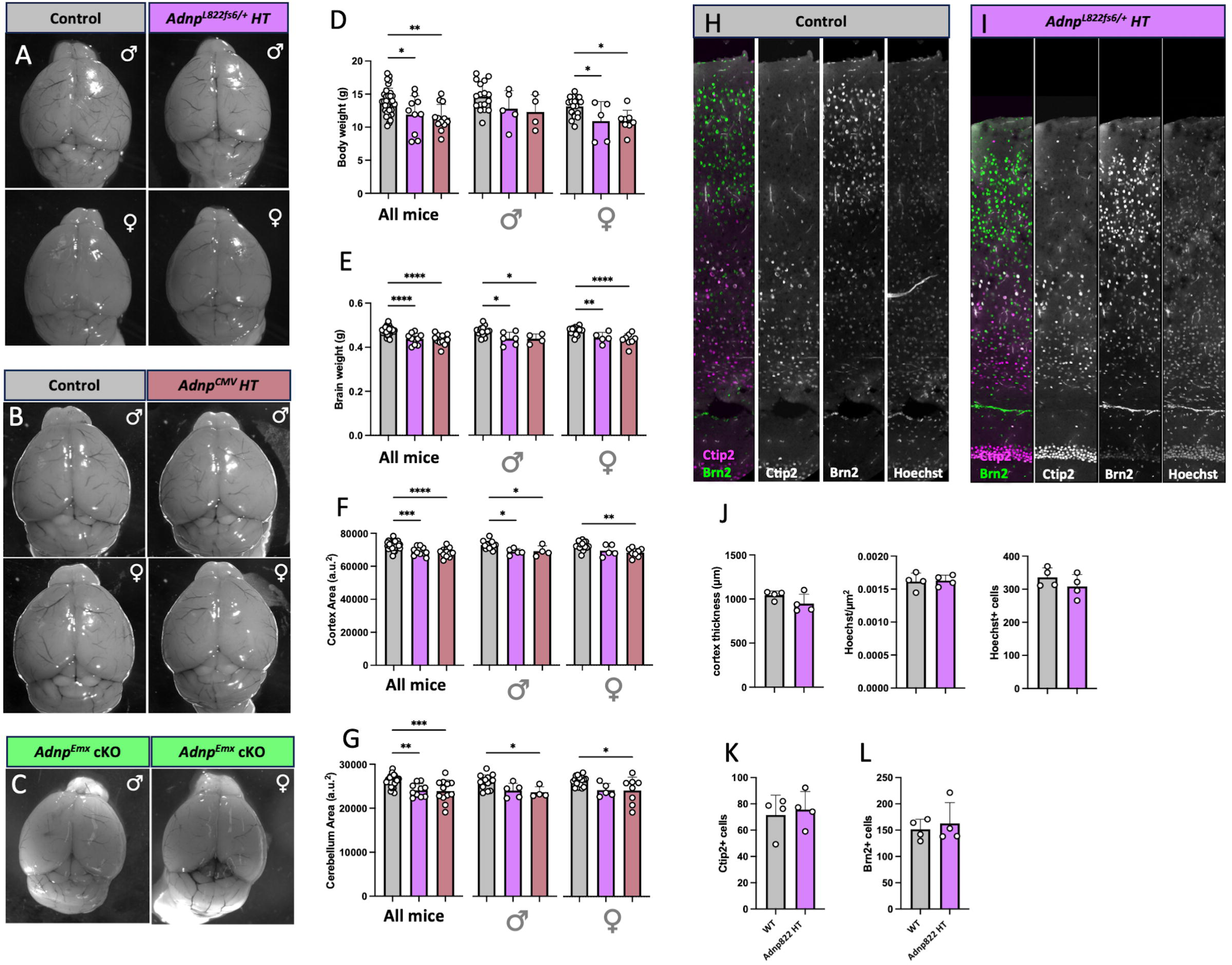
Comparison of *Adnp* mutant brains. (A-C) Control vs. *Adnp^L822fs6/+^* (A), *Adnp^Del/+^*(B) gHet brains, or *Adnp^Emx^* cKOs (C). (D) Brain weight measures. (E) Body weight measures, (F) Cortex area measures, and (G) cerebellar area measures (arbitrary units) of control, *Adnp^L822fs6/+^*gHet, and *Adnp^Del/+^* gHet brains. (H, I) Immunofluorescence staining on control (H) or *Adnp^L822fs6/+^* (I) somatosensory cortex for the lower-layer marker Ctip2 (Bcl11b) and the upper-layer marker Brn2 (Pou3f2). Hoechst counterstaining marks DNA. (J) Measurements of cortex thickness, Hoechst+ cell density, and overall cell number (200 μm-wide bins) as indicated. (K, L) Number of Ctip2+ cells (K) or Brn2+ cells (L) counted from 200 μm-wide bins. \**p*□<□0.05; \*\**p*□<□0.01; *** *p*□<□0.001; \*\*\*\**p*□<□0.0001.

### Adnp^Emx^ cKOs exhibit increased anxiety and/or exploratory behaviours

We next tested for potential changes in behaviors across mouse models. Of note, all testing was performed blinded to genotype. In terms of gross anatomy (Fig. 2) and parallel behavioral work, we noted that *Adnp^Del/+^* and *Adnp^L822fs6/+^*exhibited indistinguishable characteristics. We therefore focused on comparing behavioral phenotypes between *Adnp^L822fs6/+^* gHets versus *Adnp^Emx^* cKOs, as well as *Adnp^Flox/+^* cHet littermates.

We began by examining circadian activity patterns use the ‘beam break’ apparatus (see methods). Though *Adnp^Emx^* cKOs exhibited small decreases in ambulatory activity relative to controls (Fig. S3Aii), all *Adnp* mutants exhibited similar baseline ambulatory and total activity trends, suggesting that circadian rhythms of *Adnp* mutants were largely unaffected (Fig. S3A, B).

Next, we assessed exploratory behaviors. In the elevated plus maze test (EPM), animals were allowed to explore a ‘plus’-shaped maze with open and closed arms under low-light (100 lux) conditions (Fig. 3A). *Adnp^Emx^* cKOs exhibited increased frequency and duration in the open arms (Fig. 3B), decreased duration in the closed arms (Fig. 3C), and a slight increase in velocity throughout testing (Fig. 3D, Fig. S4G). These findings suggest that *Adnp^Emx^* cKOs may have reduced aversion to the open arms and/or increased propensity to explore the test arena. During the open field (OF) test (Fig. 3E), mice were placed in the middle of an enclosed field under a centered bright light (300 lux). *Adnp^Emx^* cKOs spent more time in the central zones relative to controls (Fig. 3Fi), though when data were separated by sex, this increase was found to reach significance only in males (Fig. 3Fii, iii). Further examination revealed that *Adnp^Emx^* cKOs had increased latency to the corners (Fig. 3G) and decreased velocity (Fig. 3K). Examination of videos (not shown) and binned time intervals (Fig. S4H-I) confirmed that *Adnp^Emx^* cKOs ‘froze’ at the center of the open field at the beginning of the test, as opposed to returning to the center multiple times. This pattern of behavior might reflect elevated anxiety within the *Adnp^Emx^* cKOs. Interestingly, male *Adnp^Emx^* cKOs exhibited normal velocity after the first minute of testing (Fig 3Hii, Fig. S4Hii), while female *Adnp^Emx^* cKOs maintained decreased velocity throughout the test (Fig 3Hiii, Fig. S4Hiii). Interestingly, we previously observed an identical ‘freezing’ behavior in *Chd4^Emx^* cKOs (Larrigan et al., 2023), suggesting that this is a ChAHP -specific phenotype.

**Figure 3.**
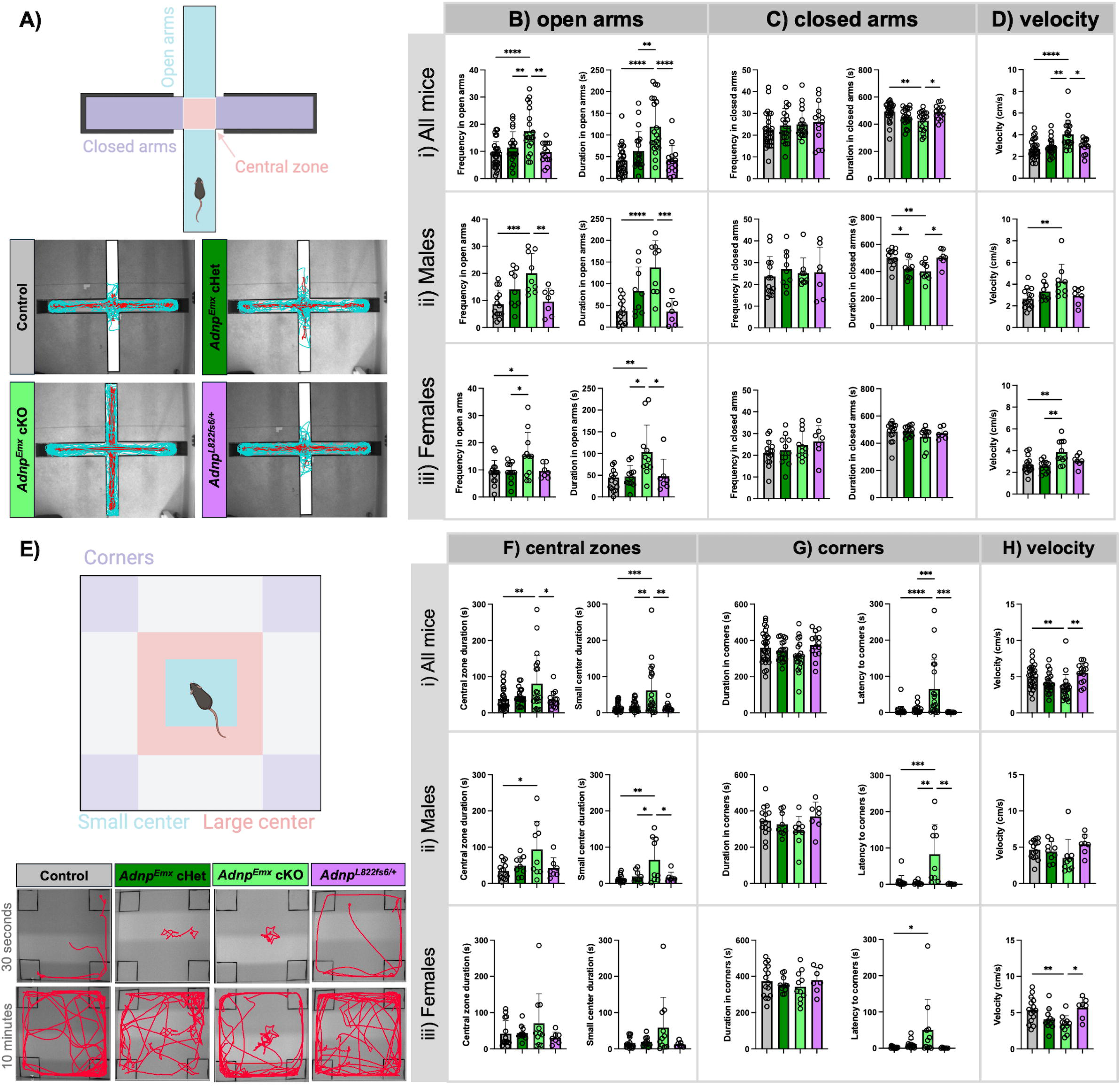
Altered exploratory and anxiety-related behaviors in *Adnp^Emx^* cKOs. (A) Schematic of elevated plus maze with two open arms and two enclosed arms, and examples of individual traces. (B-D) Frequency and duration entering open arms (B) closed arms (C), and velocity (D). (E) Schematic of open field test demonstrating boundaries for the small central zone, large central zone, and corners, and examples of individual traces. (F) Total duration in the central zone and (left) and small central zone (right). (G) Total duration in corners (left) and latency of mice to reach the corners (right) after being placed in the center of the open field. (H) Velocity during elevated plus maze. Data for control (n=29 total, n=14 males, 15 females), *Adnp^Emx^* cHets (n=20 total, n=9 males, 11 females), *Adnp^Emx^* cKOs (n=20 total, n=9 males, n=11 females) and *Adnp^L822fs6/+^* gHets (n=14 total, n=7 males, 7 females) groups are plotted showing data for all mice (i), male mice only (ii), and female mice only (iii). Statistical analyses are via one-way ANOVA with the Tukey-Kramer post-hoc test. \**p*□<□0.05; \*\**p*□<□0.01; *** *p*□<□0.001; \*\*\*\**p*□<□0.0001.

*Adnp^L822fs6/+^* and *Adnp^Emx^* cHets appeared largely unaffected in both the EPM and OF, with the exception of male *Adnp^Emx^*cHets who exhibited increased duration within the central zone (Fig. S4Dii) and decreased duration within the closed arms in EPM (Fig. S4Fii). Thus, while male *Adnp^Emx^* cHets spend more time with their noses in the central zones, they did not venture more into the open arms like their male *Adnp^Emx^*cKO counterparts. Taken together, these results suggest that Adnp plays an important role within the *Emx1-Cre* lineage that can directly affect exploratory and/or anxiety-like behavior.

### Mild effects on repetitive movements and social behavior

The marble burying test reflects repetitive behaviors and often reveals phenotypes in mouse models of autism spectrum disorder. However, we observed that male and female *Adnp^Emx^* cKOs buried none of the marbles, resulting in a significant decrease relative to controls (Fig. 4B). This may suggest aversion to interacting with the marbles, or perhaps disinterest in novel objects. Unlike the EPM and OF tests, *Adnp^Emx^* cKOs exhibited no changes in velocity (Fig. S5A). *Adnp^L822fs6/+^* and *Adnp^Emx^* cHets exhibited no changes relative to controls (Fig. 4B).

**Figure 4.**
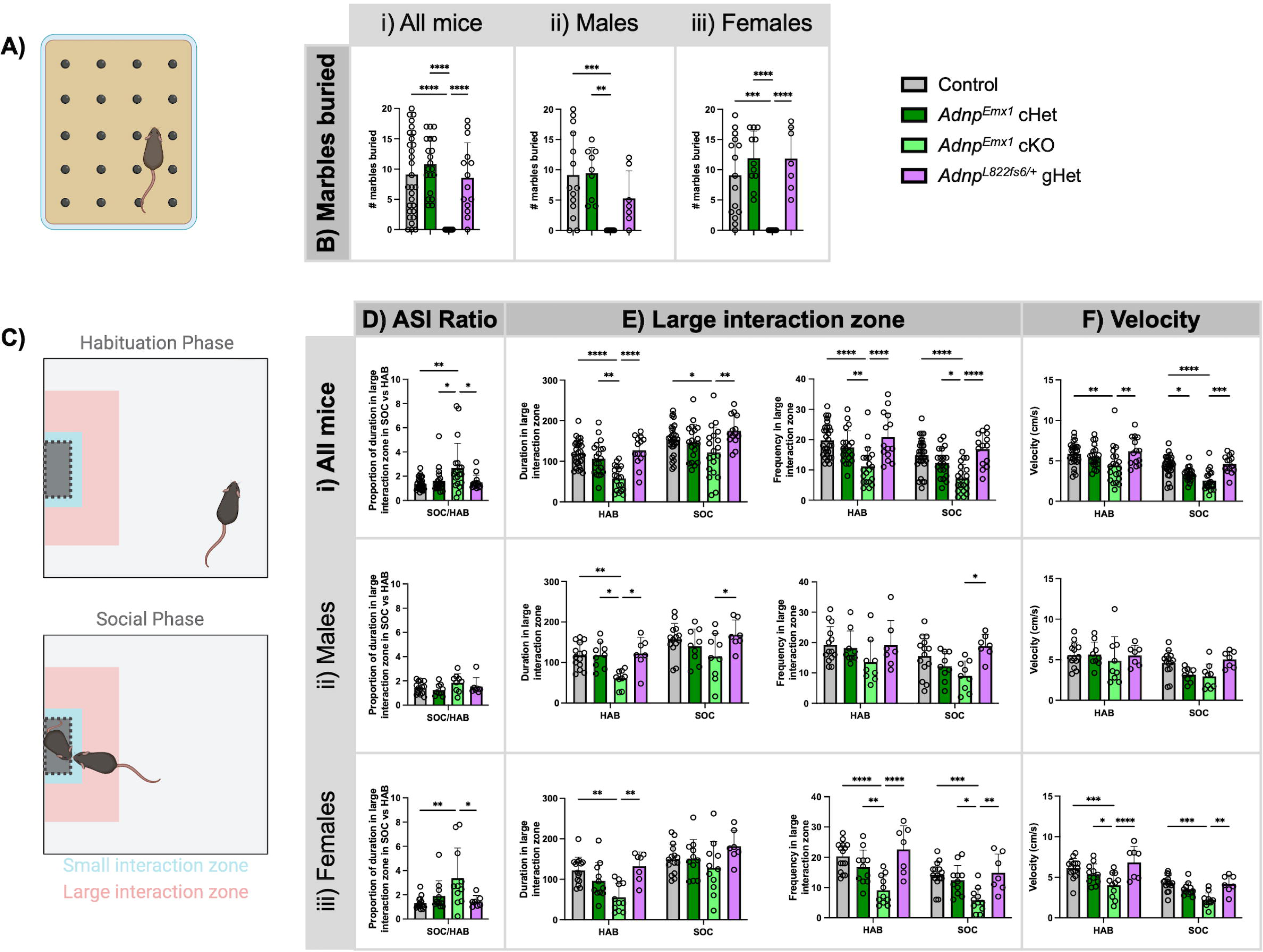
Mild phenotypes in marble burying and adult social interaction tests. (A) Schematic of marble burying test. (B) Number of marbles buried during testing. (C) Schematics of the habituation phase (no partner mouse present) and social phase (partner mouse present) for the adult social interaction test. (D) Ratio of time spent in the large interaction zone duration during the social phase relative to the habituation phase. (E) Total duration in large interaction zone (left) and frequency entering large interaction zone (right). (F) Velocity during testing. Data for control (n=29 total, n=14 males, 15 females), *Adnp^Emx^* cHets (n=20 total, n=9 males, 11 females), *Adnp^Emx^* cKOs (n=20 total, n=9 males, n=11 females) and *Adnp^L822fs6/+^* gHets (n=14 total, n=7 males, 7 females) groups are plotted showing data for all mice (i), male mice only (ii), and female mice only (iii). Statistical analyses are via one-way ANOVA with the Tukey-Kramer post-hoc test for panels B and D. Statistical analyses are via two-way ANOVA with the Tukey-Kramer post-hoc test for panels E and F. \**p*□<□0.05; \*\**p*□<□0.01; *** *p*□<□0.001; \*\*\*\**p*□<□0.0001.

*Adnp^Emx^* cKOs similarly exhibited subtle changes in behavior during the adult social interaction test (Fig. 4C). *Adnp^Emx^* cKOs increased the proportion of time spent in the large interaction zone during the social phase of the test relative to the habituation phase (Fig 4Di), though this difference was only significant in females when data was separated by sex (Fig 4Diii). Further examination revealed that *Adnp^Emx^* cKOs spent significantly less time in the large interaction zone during habituation (Fig. 4Eii, iii), which likely indicates that the phenotype is not purely driven by social context. Notably, though *Adnp^Emx^* cKOs exhibited decreased velocity during both phases (Fig. 4F), *Adnp^Emx^*cKOs only exhibited decreased velocity within the first minute of testing during the habituation phase (Fig. 4F, Fig. S5B), reminiscent of the ‘freezing’ behavior observed at the outset of OF testing. In contrast, decreased velocity was maintained throughout the social phase (Fig S5C). These effects remained significant in females when data were separated by sex (Fig 4Eii, Fig. S5Bii, Cii). This suggests that *Adnp^Emx^* cKOs display subtle changes in behavior (i.e. reduced velocity) in the presence of social partner mice.

While *Adnp^Emx^* cHets were equivalent to wild-type controls in social behavior, male *Adnp^Emx^* cHets spent more time in the small interaction zone (Fig. S5D) and entered the small interaction zone with increased frequency during habituation (Fig. S5E), perhaps suggesting increased interest in novel objects, like the empty chamber that houses social partner mice.

### Adnp^Emx^ cKOs exhibit motor deficits

Rotarod testing was performed over the course of four days to further assess sensorimotor activity and learning. *Adnp^Emx^* cKOs had decreased latency to falling compared to controls (Fig. S6A) on days 2-4 of testing, which may be indicative of a motor, learning or motivation phenotype. Interestingly, *Adnp^L822fs6/+^* exhibited increased latency to fall relative to controls on the first day of testing (Fig. S6i, Ci), which remained significant in females when data was separated by sex (Fig. S6Aiii, Ciii).

### Adnp^Emx1^ cKOs and Adnp^L822fs6/+^ gHets exhibit similar sex specific changes in learning

The Morris Water maze (MWM) is a classic test of spatial learning and memory, in which mice must learn find a submerged and hidden platform across repeated trials using visual cues (Fig. 5A). During days 3-5 of training, *Adnp^Emx^* cKOs exhibited increased latency to reach the platform (Fig. 5Bii, Biii). On day 6 of training, male *Adnp^Emx^* cKOs were able to locate the platform comparably versus controls (Fig. 5Bii), while female *Adnp^Emx^* cKOs maintained increased latency (Fig. 5Biii). During probe day, when the platform was removed to assess memory of the platform location, *Adnp^Emx^* cKOs entered the platform area significantly less times than controls (Fig. 5Ci), though this difference was lost when data was separated by sex. *Adnp^Emx^* cKOs also exhibited decreased latency during reversal training days 2-3 (Fig. 5Ei), which was maintained in female *Adnp^Emx^* cKOs when data was separated by sex (Fig. 5Eiii). Additionally, female *Adnp^Emx^* cKOs exhibited increased thigmotaxis during probe day (Fig. 5Ci, Ciii) and reversal probe day (Fig. 5Fi, Fiii), which can reflect elevated anxiety, fear and/or stress.

**Figure 5.**
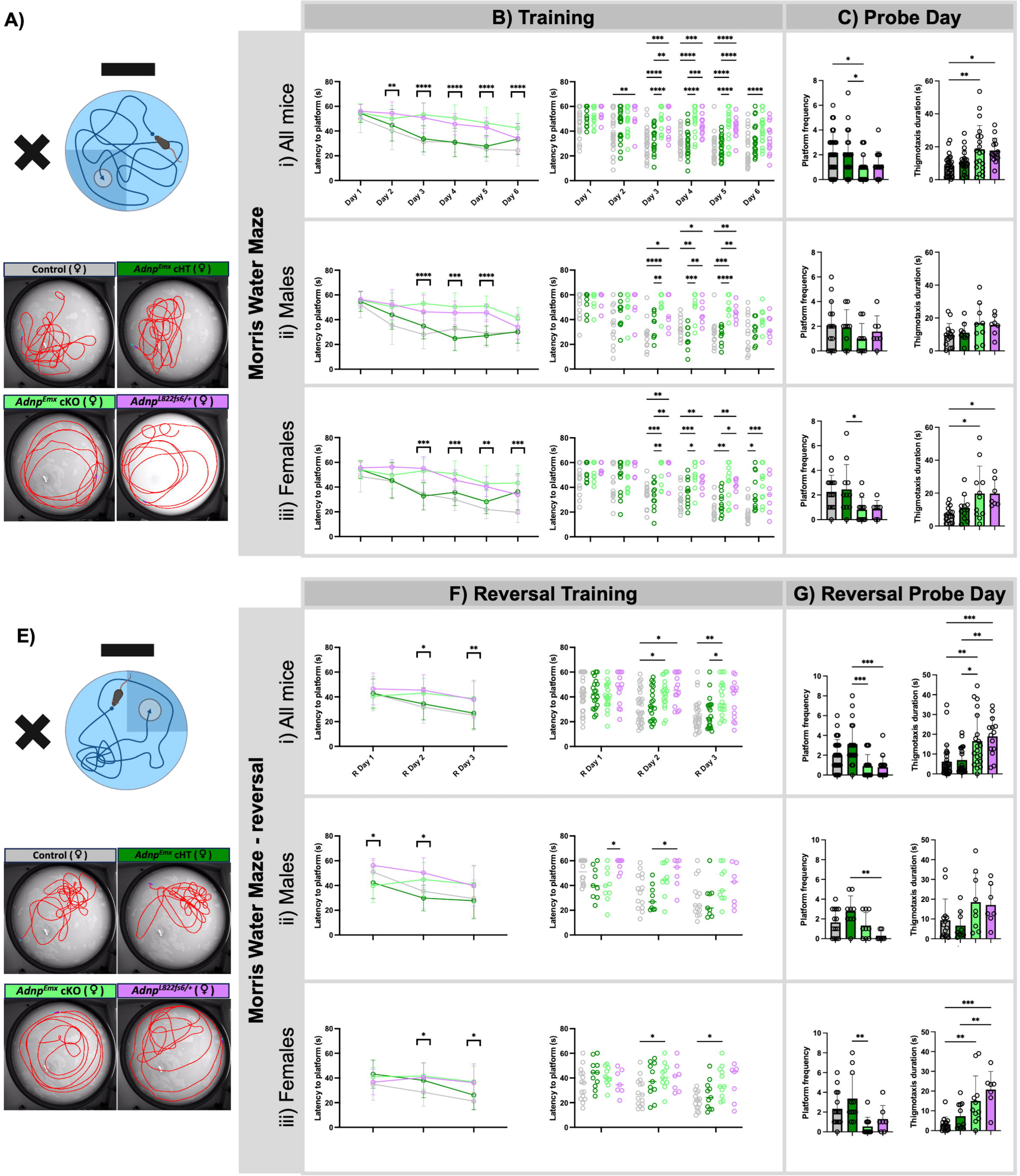
Sex-specific phenotypes in learning during the Morris Water Maze test are robust to different dosages. (A) Schematic of Morris Water Maze test (top) with labelled zones (bottom), and examples of individual traces. (B) Latency to reach hidden platform during days 1-6 of training. (C) Frequency crossing platform zone after platform is removed during probe day (left), and thigmotaxis duration during probe day (right). (D) Schematic of reversal Morris Water Maze test (top) with labelled zones (bottom). (E) Latency to reach hidden platform during days 1-6 of reversal training. (F) Frequency crossing platform zone after platform is removed during reversal probe day (left), and thigmotaxis duration during reversal probe day (right). Data for control (n=29 total, n=14 males, 15 females), *Adnp^Emx^* cHets (n=20 total, n=9 males, 11 females), *Adnp^Emx^* cKOs (n=20 total, n=9 males, n=11 females) and *Adnp^L822fs6/+^* gHets (n=14 total, n=7 males, 7 females) groups are plotted showing data for all mice (i), male mice only (ii), and female mice only (iii). Statistical analyses are via one-way ANOVA with the Tukey-Kramer post-hoc test for panels B and E. Statistical analyses are via two-way ANOVA with the Tukey-Kramer post-hoc test for panels C and F. \**p*□<□0.05; \*\**p*□<□0.01; *** *p*□<□0.001; \*\*\*\**p*□<□0.0001.

*Adnp^Emx^* cHets appeared to exhibit no change in behavior in the MWM relative to controls. However, male and female *Adnp^L822fs6/+^*heterozygotes exhibited increased latency to reach the platform on days 2-5 of training (Fig. 5B), as well as during day 2 of reversal training (Fig. 5E), though the latter change was lost when data was separated by sex. Since Rotarod testing suggested a potential motor impairment in *Adnp^Emx^* cKOs, we verified that velocity of the mice on probe and reversal probe day was not significantly affected (Fig. S6A, Fig. S7A). However, further examination on reversal probe day revealed that *Adnp^L822fs6/+^* gHets spent significantly more time in the platform’s prior quadrant versus the reversal quadrant (Fig. S7B), suggesting that *Adnp^L822fs6/+^* gHets successfully learned the platform’s location despite needing more days of training to do so. These results may also suggest inflexibility in learning the new platform location. Interestingly, female *Adnp^L822fs6/+^* heterozygotes exhibited increased thigmotaxis on probe day and reversal probe day. Thus, *Adnp^L822fs6/+^*phenotypes closely resemble *Adnp^Emx^* cKOs despite differences in Adnp dosage.

After completion of the MWM, we ran a visual test wherein a visible flag is placed above the platform, while the other visual cues are removed. During this test, female *Adnp^Emx^* cKOs had an increased latency to the platform (Fig. S7E), which may suggest impaired vision. To clarify this finding, we ran an optomotor test to better assess visual acuity. All mice exhibited similar optomotor reflexes, suggesting that the results in MWM are not due to impaired vision (Fig. S7F).

Next, we examined an aversive conditioning paradigm. During fear conditioning training (Fig. S8A), mice were placed in an apparatus that was novel to them and received three tone-shock pairings. *Adnp^Emx^* cKOs displayed increased inactivity after the tone relative to controls (Fig. S8B). No significant differences were observed during context testing (i.e. re-entering the testing arena; Fig. S8C). However, during cue association testing, *Adnp^Emx^* cKOs exhibited elevated inactivity after the tone relative to controls (Fig. S8D). However, *Adnp^Emx^* cKOs were also more inactive prior to the cue relative to controls, again suggesting increased fear, anxiety, and freezing behaviors Fig. S8D). Data separated by sex suggest that these phenotypes may be more elevated in females.

## Discussion

While great strides have been made in the development of NDD model systems - including brain organoids that can be generated from the cells of affected individuals, mouse models remain a key system for bridging genetics to behavioral phenotypes. Standardized tests have been used to investigate a host of NDD mouse models, allowing many laboratories to easily replicate and extend them as bioassays for fundamental or preclinical investigation. Moreover, conditional genetics affords the opportunity to map phenotypes to specific cell types and circuits. In this regard, the six-layered neocortical circuit architectures that integrate sensorimotor information with higher order cognitive functions are only found in mammals.

Here, to help establish a baseline for such investigations, and to explore questions of *Adnp* gene dosage, we generated and compared an allelic series of germline versus conditional genetic mouse models. By using a blinded and largely automated testing approach, we were able to identify neuroanatomical and behavioral phenotypes shared between genotypes irrespective of dosage. Since we focused on mouse lines that have already been shared with repositories for public distribution (Clemot-Dupont et al., 2025; D’Incal et al., 2026), we hope that this work will provide a useful resource for the broader NDD research community.

### An allelic series underscores the link between Adnp, cortical neurogenesis, and viability

To first assess the requirement for *Adnp* throughout the neural tube, we generated *Adnp^Nestin^* cKOs. While, *Adnp^Nestin^*cKOs could not be recovered postnatally, Mendelian frequencies were observed at E15.5. Although germline *Adnp* knockouts are also non-viable (Pinhasov et al., 2003), germline *Adnp* homozygotes exhibit defects in exiting pluripotency and fail to generate neural progenitors – which is also seen when *ADNP* is deleted in pluripotent cells (Ostapcuk et al., 2018; Sun et al., 2020a; Yan et al., 2022). Since the neural tube is generated prior to *Nestin-Cre* onset, we surmise that *Adnp^Nestin^* cKOs likely die due to a completely different mechanism. Indeed, we found that *Adnp^Nestin^*cKOs exhibited deficits in the production of upper layer neurons without reductions in early-born Bcl11b+ lower-layer neurons. This correlated well with our previous observations in the *Adnp^Emx^* cKO (Clemot-Dupont et al., 2025), and these cKOs also closely match *Chd4^Nestin^* and *Chd4^Emx^* cKOs (Nitarska et al., 2016; Larrigan et al., 2023). In the neocortex, *Adnp* expression was previously shown to be elevated in upper-layer neurons and suppressed by the lower-layer determinant *Tbr1* (Notwell et al., 2016), which is itself a prominent NDD risk gene. These findings suggest a specific connection between Adnp and upper-layer neuronal specification. However, phenotypes might alternatively reflect a progressive increase in gene misregulation that reaches a phenotypic threshold only after several days, disproportionately affecting later-born cells. Nonetheless, germline and *Adnp^Nestin^* cKOs both reinforce the general link between Adnp, neurodevelopment, and viability.

Next, we examined *Adnp^Del/+^* and *Adnp^L822fs6/+^*gHets. Both models exhibited indistinguishable deficits in brain size, consistent with the notion that the *Adnp^L822fs6^* variant may act in a loss-of-function manner. Contrasting with cKO phenotypes, cell-type composition in *Adnp^L822fs6/+^* gHets appeared to be largely normal despite presenting with slight decreases in body weight, brain weight, and brain size, in agreement with prior work (D’Incal et al., 2026). Taken together, these data suggest that hypoplasia is a core phenotypic feature but varies in magnitude according to dosage, whereas changes in cell type composition emerge only when *Adnp* is fully ablated.

In agreement with prior analyses of gross anatomy and gene expression (Clemot-Dupont et al., 2025), we observed little difference between *Adnp^Emx^*cHets versus wild-type controls, though the EPM and ASI tests revealed elevations in exploratory behaviors as well as interactions with novel objects. Moreover, while *Adnp* gHets and cKOs exhibited deficits in the MWM, *Adnp^Emx^*cHets trended oppositely, with significant improvements in performance versus mutant genotypes. Interestingly, the EPM test was also the only test that appeared to have a dosage-like effect, with *Adnp^Emx^* cHets having increased duration of their nose point in the central zone, whereas *Adnp^Emx^*cKOs ventured out much more into the open arms. Curiously, the *Adnp^L822fs6/+^*heterozygotes lacked these phenotypes. One potential explanation may be that the conditional genetics approach imbalances cortical circuits since the genetic modification is restricted to the *Emx1* lineage, while other cells that contribute to circuits are unaffected - such as cortical inhibitory neurons. Alternatively, these results might reflect a functional difference between the null versus frameshifting mutations. In summary, *Adnp* phenotypes aligned with the severity predicted by genetic dosage (Fig. 6).

**Fig. 6.**
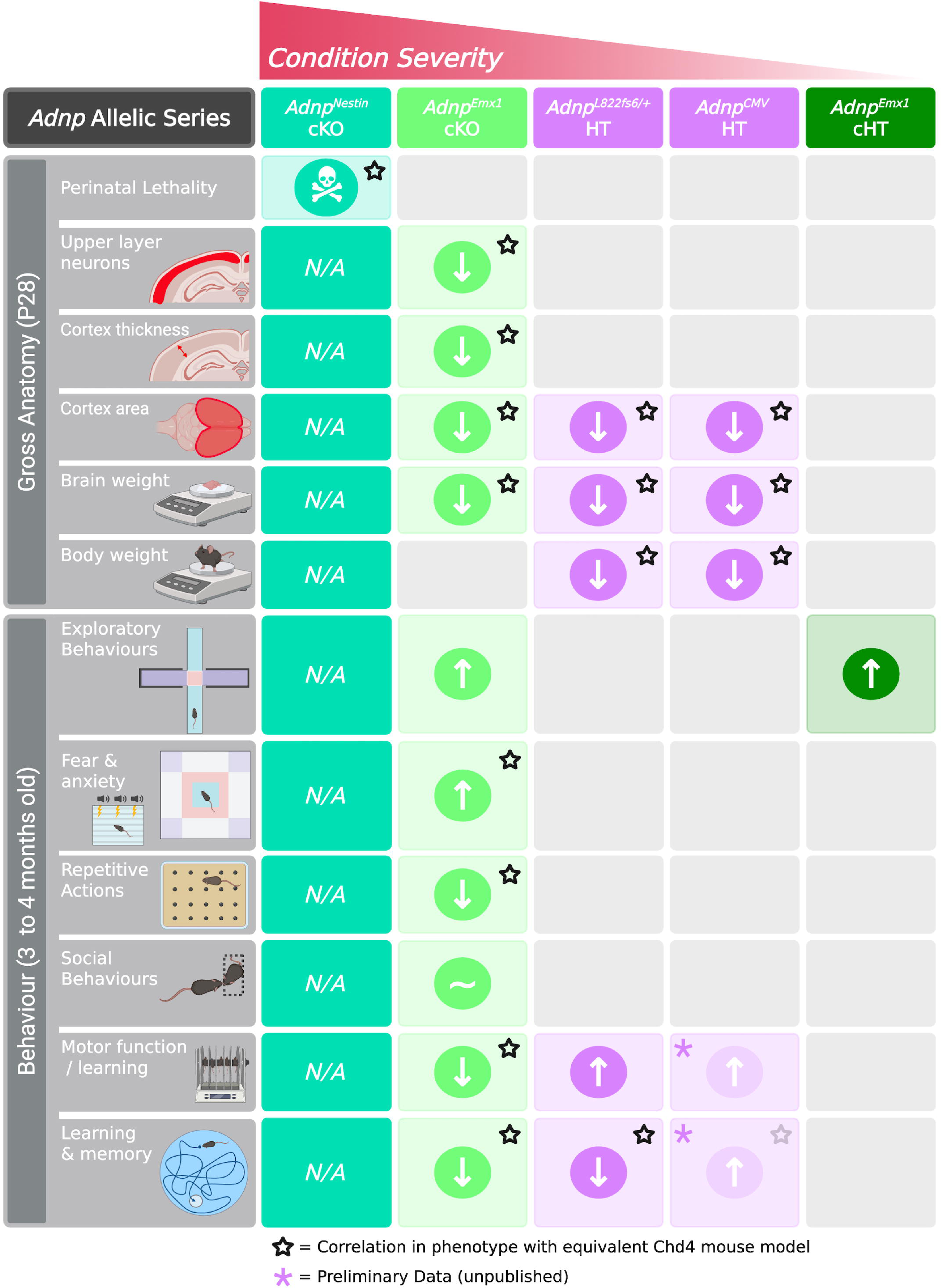
*Adnp* genotypes ranked by phenotypic severity. Phenotypic outcomes include viability, body and brain size measurements, exploratory behaviors, anxiety, repetitive behaviors, motor function, and learning. Stars indicate matched phenotypes with *Chd4^Nestin^* cKOs (Nitarska et al., 2016), as well as *Chd4^Emx^*cKOs and *Chd4^Del/+^* gHets (Larrigan et al., 2023). *Adnp^Emx^*cHet and cKO neuroanatomical measures were previously reported (Clemot-Dupont et al., 2025). Comparative *Adnp^Del/+^* gHet behavioral data is preliminary and will be published elsewhere.

### Adnp in learning and anxiety

To identify core behavioral features that manifest across the allelic series, we focused on *Adnp^Emx^* cKOs and cHets, as well as *Adnp^L822fs6/+^* gHets. Strikingly, *Adnp^Emx^* cKOs and *Adnp^L822fs6/+^* gHets exhibited indistinguishable phenotypes in learning during the MWM test, and even shared sex-specific phenotypic features. Males displayed increased latency until the final day of testing, and females exhibited increased latency throughout testing and increased thigmotaxis on probe day. In contrast, these phenotypes were not observed in *Adnp^Emx^* cHets. This suggests that *Adnp* haploinsufficiency specifically within the *Emx1* lineage is not sufficient to undermine learning and memory. While this result might imply that *Adnp^Emx^* cKOs and *Adnp^L822fs6/+^* gHets manifest convergent phenotypes via distinct mechanisms that might involve additional circuits, the MWM test depends on neocortical and hippocampal circuits that are captured within the *Emx1 lineage*. One potential explanation is that germline heterozygosity reduces Adnp dosage much earlier, and because *Emx1-Cre* initiates much later, that full ablation is required to generate a comparable genetic insult. Alternatively, since *Adnp^Emx^* cHets do not exhibit detectable growth defects, whereas germline heterozygotes and cKOs have hypoplastic brains, the behavioral deficits might be tied to neuroanatomical alterations – even though *Adnp^L822fs6/+^* gHets did not exhibit the drastic shifts in cortical size and cell-type composition exhibited by the cKOs (Clemot-Dupont et al., 2025).

The observed thigmotaxic phenotypes are suggestive of elevated anxiety, fear and/or stress in mice, and were evident in both *Adnp^Emx^*cKOs and *Adnp^L822fs6/+^* female gHets during MWM testing. However, data from other tests suggests that anxiety-like phenotypes are much more prevalent in *Adnp^Emx^* cKOs. Most notably, *Adnp^Emx^*cKOs displayed decreased velocity across tests, including the OF and FC tests assessing for fear and anxiety. Males initially froze longer and then resumed normal behavior, whereas females initially froze for shorter periods of time but maintained decreased velocity throughout the OF test. Females also specifically maintained reduced velocity in ASI testing in the presence of social mice. Conversely, BBK testing suggested that male *Adnp^Emx^* cKOs have larger reductions in spontaneous ambulatory activity. While most of these phenotypes were not found in cHets or gHets, elevated thigmotaxis was observed in *Adnp^L822fs6/+^* females.

Elevated anxiety might be causally linked to the observed neuroanatomical differences. Our prior study links Adnp to cortical growth, but also suggests that Adnp is particularly important for gene expression programs associated with upper-layer neurons (Clemot-Dupont et al., 2025). While upper-layer neurons were not reduced in number in *Adnp^L822fs6/+^* gHets, re-analysis of published RNA-seq data (D’Incal et al., 2026), suggests that key transcription factors associated with upper-layer identity are systematically downregulated in *Adnp^L822fs6/+^* gHets, while lower-layer markers are generally unaffected. We think that imbalances in neuron subtype specification could contribute towards the observed anxiety-like phenotypes. Indeed, the amygdala, which plays an important role in fear and anxiety, has been shown to project to neurons within the prelimbic and infralimbic cortex, both which fall within the *Emx1* lineage. Moreover, regions of the amygdala itself also fall within the *Emx1* lineage (Gorski et al., 2002; Cheriyan et al., 2016). However, while *Adnp^Emx1^* cKOs displayed impairments in learning during the MWM test, the fear conditioning test was not affected – despite the fact that this form of aversive learning depends on input from the limbic system. Future work will be required to determine how exactly Adnp modifies circuits associated with learning and anxiety.

### Behavioral profiling links the ChAHP complex to learning

Notably, many of the behaviors exhibited by *Adnp* models closely matched the phenotypes that we previously observed when analogous *Chd4* models were tested using an identical workflow (Larrigan et al., 2023). For example, both *Adnp^Emx^* and *Chd4^Emx^* cKOs (but not gHets) appeared to freeze in the center of the arena at the onset of OF test. In the marble burying test, both *Adnp* and *Chd4* cKOs (but not gHets) generally failed to bury any marbles. Both of these behavioral manifestations are highly atypical, and we think they may reflect heightened anxiety. Moreover, it is striking that these captured *Adnp^Emx^* cKO phenotypes corresponded most closely with the parallel *Chd4^Emx^* cKO model - rather than *Adnp* models with different genetic dosages.

Importantly, for both *Adnp* (this study) and *Chd4* (Larrigan et al., 2023), the MWM was the only test in which gHets and cKOs exhibited quantitatively similar deficits. Taken together, our data imply that the ChAHP complex is responsible for many of the key anxiety and learning behavioral phenotypes exhibited by *Adnp* mutants (Fig. 6; Fig. S9). Accordingly, Chd4 is Adnp’s most significant protein partner in unbiased proteomic studies (Ostapcuk et al., 2018; Sharifi Tabar et al., 2022; Pintacuda et al., 2023). However, a few phenotypes were less well aligned. For example, *Chd4^Emx^* cKOs and cHets displayed no changes in exploration in the EPM test (Larrigan et al., 2023). Moreover, prior work focused on the cerebellum suggested that Chd4 can regulate behavior via the NuRD complex (Yang et al., 2016), which is distinct from ChAHP (Ostapcuk et al., 2018). These observations might reflect ChAHP-independent functions that could underpin phenotypic divergence between HVDAS and SIHIWES.

Behavior emerges from numerous circuits and processes that are challenging to dissect experimentally. It will be important to further investigate and understand these processes, and we think that direct comparison between *Adnp* and *Chd4* mutant models is likely warranted. While our prior work showed that ChAHP exerts a strong influence on the regulation of many NDD risk genes during embryonic development (Clemot-Dupont et al., 2025), we have little mechanistic understanding of how ChAHP remodels chromatin in the adult brain. It will therefore be critical to study how ChAHP regulates gene expression in mature neurons and glia in the future.

## Methods

### Animal work

All animal work was approved by the uOttawa Animal Care Committee and carried out under ethical protocols OHRI-2856, OHRI-3155, OHRI-3949, OHRI-4029, and OHRI-4178, following guidelines set out by the Canadian Council of Animal Care. Animals were housed in the *uOttawa Animal Care and Veterinary Services* facility under a 12-h light cycle with food and water provided *ad libitum*. The *Adnp^Flox^* allele was previously described (Clemot-Dupont et al., 2025), and has been deposited in the Jackson Laboratory (*B6(Cg)-Adnp^em1Pmtr^/J*, Stock no. 039849). *Nestin-Cre+* mice (Berube et al., 2005) and *Emx1-Cre+* mice (Gorski et al., 2002) were generously provided by the David Picketts laboratory (OHRI). *CMV-Cre* mice (Schwenk et al., 1995) were purchased directly from Jackson Labs (*B6.C-Tg(CMV-cre)1Cgn/J*, Stock no. 006054). All mice were backcrossed to C57BL/6J background mice for at least six generations before data collection. Genotyping was performed using primer sets listed in the corresponding referenced publications. See also Table S1.

### Immunofluorescence

Immunofluorescence staining was performed as previously described (Larrigan et al., 2023; Clemot-Dupont et al., 2025). Briefly, mice were anesthetized with isoflurane prior to euthanasia via anesthesia CO2 inhalation followed by cervical dislocation. After dissection, brains were weighed, imaged, and fixed in 4% PFA overnight. Brains were subsequently transferred to 20% sucrose for 24 h, freezing solution (1:1 v/v of OCT:20% sucrose in PBS) for 24-36 h, and frozen down to −80°C in freezing solution. Brains were sectioned coronally using a Leica CM1860 cryostat. Thickness was set at 12 μm for prenatal brains and 16 μm for postnatal brains. Sections were rinsed in 1× PBS followed by antigen retrieval in citrate buffer (10 mM Sodium citrate, 0.05% Tween 20, pH 6.0) using a pressure cooker for 10 min, and rinsing with water for 10 min. Primary antibodies were incubated overnight at 4°C, followed by three 1X PBS washes, incubation with secondaries for 1 h at room temperature, three 1× PBS washes and mounting in Mowiol mounting media (12% w/v Mowiol 4–88, 30% w/v glycerol, 120 mM Tris-Cl pH 8.5). All antibodies are listed in Table S2 and were diluted in 3% w/v Bovine Serum Albumin, 0.4% Triton, with Hoechst 33342 in PBS.

### In situ RNA hybridization

Section *in situ* hybridization was performed as described previously (Mattar et al., 2004; Touahri et al., 2015). A digoxigenin-labelled antisense probe was generated from a linearized pBluescript II plasmid harboring an *Adnp* ORF cDNA. The original *Adnp* cDNA was subcloned from pEntr223 *Adnp* (MmCD00295392), obtained from D.N.A.S.U. (University of Arizona).

### Cell counting and imaging

Dissected cortex and cerebellum area imaging and measurements was performed as previously described (Larrigan et al., 2023; Clemot-Dupont et al., 2025). For consistency across coronal sections, the primary somatosensory cortex located above the hippocampus was targeted for imaging and cell counting. For each slide, every other section was imaged using the Zeiss LSM900 Confocal Microscope at 200× magnification. Single Z-planes were tiled and stitched using Zen software (Zeiss). Manual cell counting was performed using the ‘Cell Counter’ Plugin in Fiji (Schindelin et al., 2012) on 200 μm-wide sections. Images of P28 cortices were separated into six 200 μm-wide bins of equal height for counting. Images of E15.5 embryonic cortices were separated into zones for counting based on histological criteria: cortical plate (CP), intermediate zone (IZ), subventricular zone (IVZ) and ventricular zone (VZ). The cortical plate was further subdivided into four bins based on Bcl11b staining: the first bin contained cells above the superficial border of Bcl11b staining, and the remaining three bins were sized to be equal in height (see Fig. S1F). The ‘Straight Line’ tool in Fiji was used to measure thickness of the cortex and cortical layers along the radial axis. Three technical replicates were counted for each biological replicate (animal) for both measurements and cell counting.

### Statistics

Data are presented as mean□±□standard deviation unless otherwise indicated. Symbols in graphs, as well as n-values in text, represent individual biological replicates (different animals). Sample sizes were not predetermined by power calculations. Statistical analysis was performed via Student’s t-test, one-way ANOVA with the Tukey/Kramer post-hoc test, or two-way repeated measures ANOVA with Tukey’s multiple comparison test as indicated in text. For all data, significance was considered as P < 0.05 and presented as ^∗^ p < 0.05, ^∗∗^ p < 0.01, ^∗∗∗^ p < 0.001, and ^∗∗∗∗^ p < 0.0001. All statistical analyses and graphs were performed/generated using GraphPad Prism version 10.6.1 for macOS, GraphPad Software, San Diego, California USA, www.graphpad.com. No data points were excluded in this study.

### Behavior

Behavioral testing was performed through the *Animal Behaviour and Physiology Core* at uOttawa as previously reported (Larrigan et al., 2023). Modifications and extensions added to the testing paradigm are described below.

### Morris water maze – Reversal

Morris water maze training and probe were performed as described previously (Larrigan et al., 2023). Following probe testing, the platform was moved to the opposite quadrant from where training was completed. Reversal training was completed over the course of 3 days, following the same protocols used for regular training. Reversal probe day was completed on the fourth day using the same protocol for regular probe day. Following this, a visual test was performed by attaching a visible flag to the platform and removing all other visual cues. Latency for mice to reach the visible flag and platform was measured 4 times at each starting location, the order of which was randomized.

### Fear conditioning

During training, mice were placed into a four-sided box-shaped apparatus for 6 minutes. After two minutes, three tone-shock pairings (30 second tone co-terminated with a two second foot 0.30 mA shock) were administered with rest intervals in between. The following day, mice were re-introduced into the same apparatus for 6 minutes without any tone shock pairings to assess context conditioning. Freezing during the first three minutes and the last three minutes was measured and compared. On the third day, mice were introduced into a novel environment (triangular-shaped, vanilla scented, covered floor, different lightning) and the tone from training was played without the shock. Freezing prior to playing and after playing the tone was measured to assess cue conditioning.

### RNA-seq and bioinformatic analyses

Analysis of published RNA-seq data (D’Incal et al., 2026) was performed as described previously (Herrera et al., 2023), except that Fastq files were mapped to the mm10 genome using Hisat2. Data were obtained from the GEO database (GSE319683). *ADNP* pathogenic variant data were obtained from the gnomAD server (Chen et al., 2024).

---

**Table S1:**
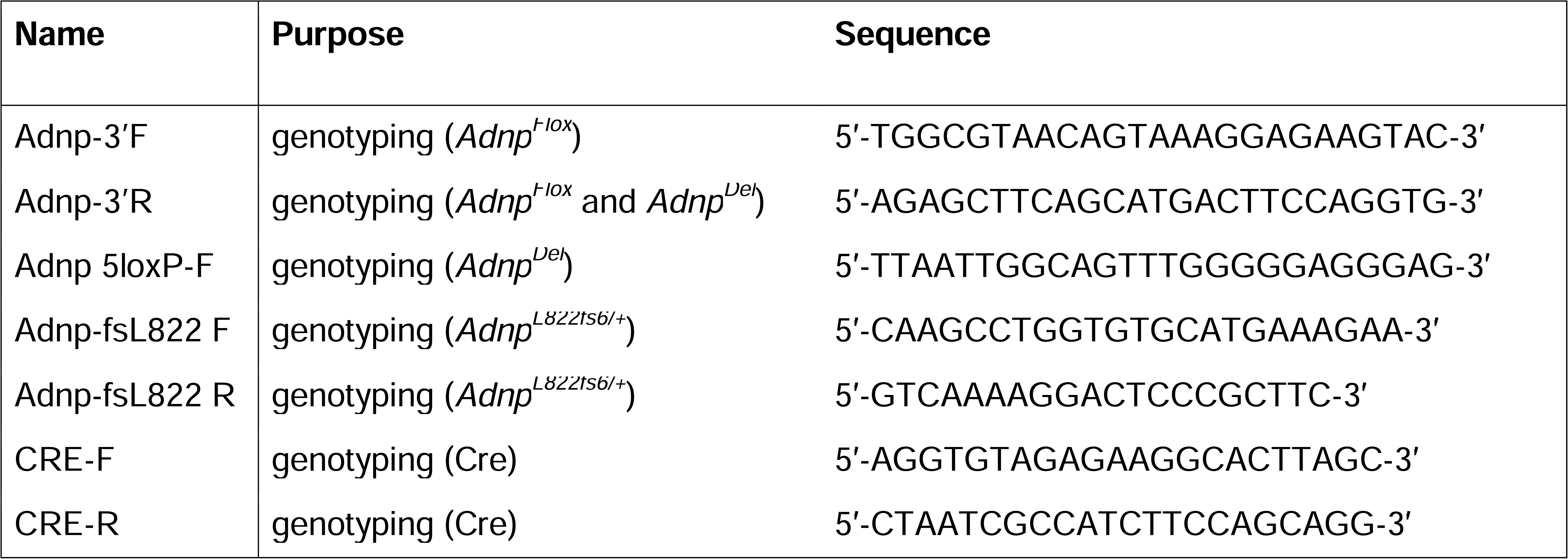
Oligonucleotide sequences.

**Table S2:**
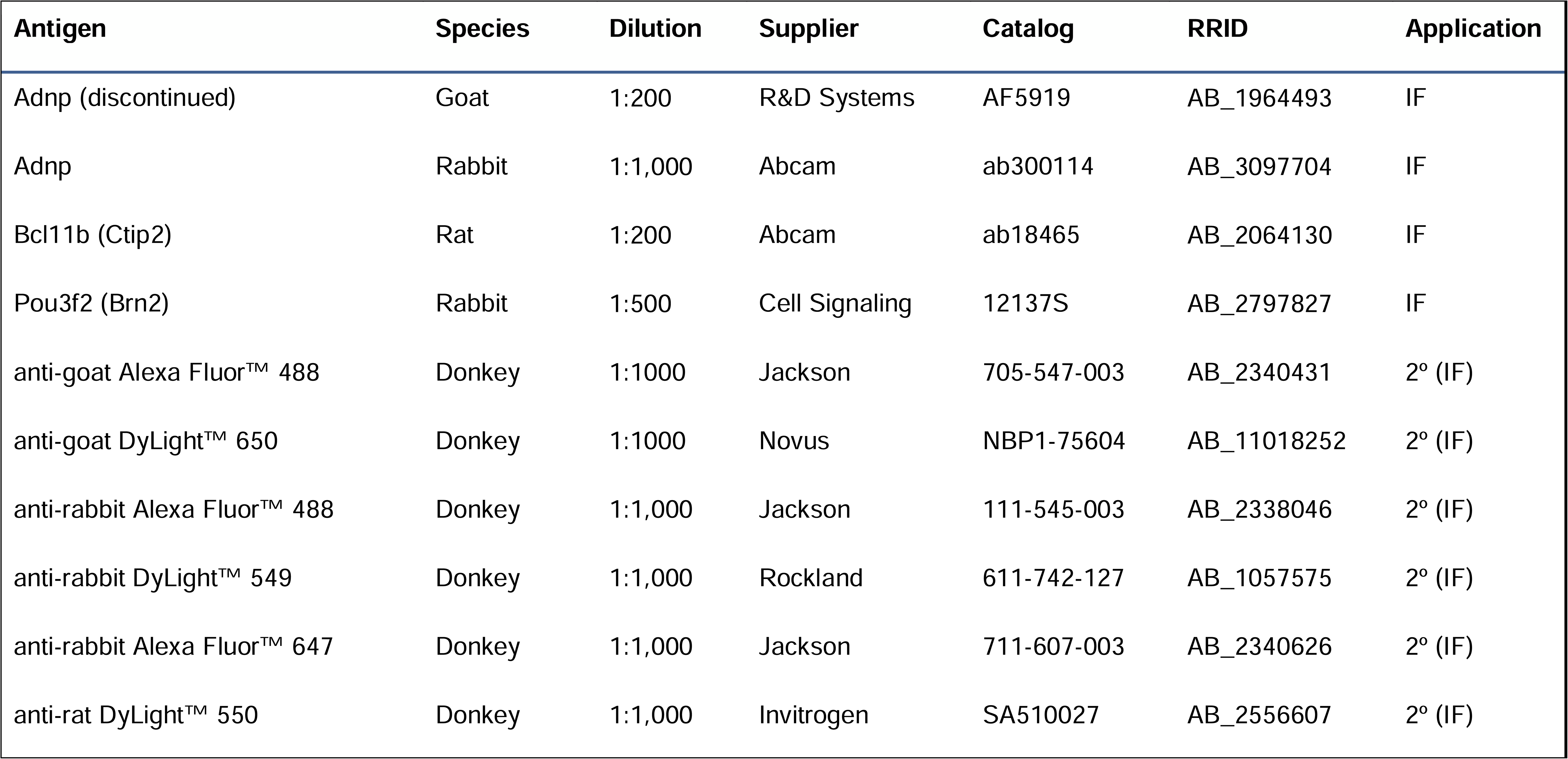
Antibodies and dilutions.

## Acknowledgements

We thank Dr. Frank Kooy for sharing mice and datasets upon which this study relied. We thank Dr. David Picketts for sharing *Emx1-cre* and *Nestin-Cre* mice. We are grateful to Kerstin Ure, Sarah Kealey, and Katherine Tacay from the uOttawa *Animal Behaviour and Physiology Core*, both for behavior testing, as well as for advice, training, and guidance. We thank Chloë van Oostende-Triplet and the *Cell Biology and Image Acquisition Core Facility*, and the staff of the uOttawa *Animal Care and Veterinary Service*. We thank Adam Baker and Rory Stanley for help and support with this project. We thank members of the Mattar and Picketts labs for their ongoing help and input. This work was generously funded by the Canadian Institutes of Health Research (CIHR) Operating Grants (PJT-166032 and OGB-198246 to P.M.). Our Adnp research program is also generously supported by the Office of the Assistant Secretary of Defense for Health Affairs through the Autism Research Program, Award No. AR240210, as well as through an Eagle’s Foundation for Autism Research grant: “Deciphering the role of chromatin remodelling complexes in ASD etiology”, awarded to P.M. and D.J.P. We additionally acknowledge the generous support of the CIHR operating grants (PJT-180577 and PJT-420505 to D.J.P.), the New Frontiers in Research Fund (NFRFT-2022-00327 to P.M.), and from the Natural Sciences and Engineering Research Council of Canada (RGPIN-2026-05722 to P.M). The project was also supported by an infrastructure grant from the Canada Foundation for Innovation for confocal microscopy (JELF 37688 to P.M.). P.M. gratefully holds the Gladys and Lorna J. Wood Chair for Research in Vision. Several figures were created in part using BioRender (https://BioRender.com).

## Author Contributions

Conceptualization: S.L., P.M. Data curation: all authors. Formal analysis: all authors. Investigation: all authors. Project administration, resources, supervision: P.M. Visualization: SL, P.M. Writing—original draft: S.L. Writing, review, and editing: all authors.

## Ethics Declarations

### Ethics Statement

Animal work was conducted according to the guidelines of the Canadian Council on Animal Care and the Animal Care and Veterinary Service at uOttawa using ethical protocols OHRI-2856, OHRI-3155, OHRI-3949, OHRI-4029, and OHRI-4178.

### Consent for Publication

All authors consent to the publication of this manuscript.

## Availability of Data and Materials

Raw behavioral data video files, data, and microscopy images are available upon request.

---

**Supplemental Figure 1. A*d*npNestin cKOs exhibit decreased upper-layer neurons at E15.5.** (A) Adnp, Pou3f2 and Bcl11b immunohistochemistry on coronal sections from E15.5 control (left) and *Adnp^Nestin^* cKO (right) brains. (B) Adnp, Pou3f2 and Bcl11b immunohistochemistry on coronal sections from E15.5 control (left) and *Adnp^Nestin^* cKO (right) cortices. (C) Thickness of cortex, (D) cell density, and (E) thickness of zones within the cortex at E15.5 for control and *Adnp^Nestin^* cKO cortices. (F) Schematic of ventricular zone (VZ), subventricular zone (SVZ), intermediate zone (IZ) and cortical plate (CP) zones and subdivided CP bins (in grey) for E15.5 cell counting. The superficial bin represents cells above the superficial Bcl11b+ border. The remaining three Bcl11b bins are equal in size. Quantification of Hoechst+ (G), Pou3f2+ (H) and Bcl11b+ (I) cells in control and *Adnp^Nestin^*cKO cortices, shown as total cells, cells per zone, and cells per CP bin. \**p*□<□0.05; \*\**p*□<□0.01.

**Supplemental Figure 2. Decreased expression of upper-layer marker genes in adult *Adnp^L822fs6/+^* gHets.** (A) Relative expression of upper-layer marker genes in control vs. *Adnp^L822fs6/+^*gHets as indicated. (B) Relative expression of lower-layer marker genes in control vs. *Adnp^L822fs6/+^* gHets as indicated. Asterisks indicate differentially regulated genes (DEGs) with adjusted p-values < 0.05 as determined via Deseq2. Data were remapped and analyzed from GSE319683 (D’Incal et al., 2026).

**Supplemental Figure 3. A*d*np mouse models exhibit similar ambulatory activities levels relative to controls.** (A) Cumulative ambulatory activity counts per hour during the beam break assay. (B) Cumulative total activity counts per hour during the beam break assay. Data for control (n=29 total, n=14 males, 15 females), *Adnp^Emx^*cHets (n=20 total, n=9 males, 11 females), *Adnp^Emx^* cKOs (n=20 total, n=9 males, n=11 females) and *Adnp^L822fs6/+^* gHets (n=14 total, n=7 males, 7 females) groups are plotted showing data for all mice (i), male mice only (ii), and female mice only (iii). Statistical analyses are via two-way ANOVA with the Tukey-Kramer post-hoc test. \**p*□<□0.05; \*\**p*□<□0.01; *** *p*□<□0.001.

**Supplemental Figure 4. Additional analyses of elevated plus maze and open field tests.** (A-G) Elevated plus maze measures, including (A) frequency in central zone, (B) duration in central zone, (C) frequency of nose point in central zone, (D) duration of nose point in central zone, (E) duration of nose point in open arms, and (F) duration of nose point in closed arms. (G) Velocity recorded in 1-minute bins. (H-K) Open field measures, including (H) velocity recorded in 1-minute bins, (I) duration in the small center recorded in 1-minute bins, (J) duration in the small center recorded in 1-minute bins for each of the individual *Adnp^Emx^*cKOs, and (K) frequency in the corners (left) and frequency in the large center (right). Data for control (n=29 total, n=14 males, 15 females), *Adnp^Emx^*cHets (n=20 total, n=9 males, 11 females), *Adnp^Emx^* cKOs (n=20 total, n=9 males, n=11 females) and *Adnp^L822fs6/+^* gHets (n=14 total, n=7 males, 7 females) groups are plotted showing data for all mice (i), male mice only (ii), and female mice only (iii). Statistical analyses are via one-way ANOVA with the Tukey-Kramer post-hoc test. \**p*□<□0.05; \*\**p*□<□0.01; *** *p*□<□0.001; \*\*\*\**p*□<□0.0001.

**Supplemental Figure 5. Additional analyses of marble burying and adult social interaction tests.** (A) Velocity during the marble burying test. (B-E) Adult social interaction measures, including (B) velocity recorded in 1-minute bins during habituation, (C) velocity recorded in 1-minute bins during the social phase, (D) duration in the small interaction zone during habituation (left) or socialization (right), and (E) frequency in the small interaction zone during habituation (left) or socialization (right). Data for control (n=29 total, n=14 males, 15 females), *Adnp^Emx^*cHets (n=20 total, n=9 males, 11 females), *Adnp^Emx^* cKOs (n=20 total, n=9 males, n=11 females) and *Adnp^L822fs6/+^* gHets (n=14 total, n=7 males, 7 females) groups are plotted showing data for all mice (i), male mice only (ii), and female mice only (iii). Statistical analyses are via one-way ANOVA with the Tukey-Kramer post-hoc test. \**p*□<□0.05; \*\**p*□<□0.01; *** *p*□<□0.001; \*\*\*\**p*□<□0.0001.

**Supplemental Figure 6. Sensorimotor phenotypes assessed via the rotarod test.** Latency to fall during the rotarod test plotted over the course of four days (A), plotted per trial (B), per day 1 trials (C), per day 2 trials (D), per day 3 trials (E), and per day 4 trials (F). Data for control (n=29 total, n=14 males, 15 females), *Adnp^Emx^*cHeTs (n=20 total, n=9 males, 11 females), *Adnp^Emx^* cKOs (n=20 total, n=9 males, n=11 females) and *Adnp^L822fs6/+^* gHets (n=14 total, n=7 males, 7 females) groups are plotted showing data for all mice (i), male mice only (ii), and female mice only (iii). Statistical analyses are via two-way ANOVA with the Tukey-Kramer post-hoc test. \**p*□<□0.05; \*\**p*□<□0.01; *** *p*□<□0.001; \*\*\*\**p*□<□0.0001.

**Supplemental Figure 7. Additional analyses of the Morris Water Maze test.** (A) Average velocity during probe day. (B) Frequency entering (right) and duration within (right) the quadrants of the Morris water maze during probe day. (C) Frequency entering (right) and duration within (right) the platform target area during probe day (Target BR), relative to the corresponding platform target areas in the other quadrants (Targets BL, FL and FR). (D) Schematic of platform reversal with example traces. (E) Latency to the platform in the Morris water maze visual test, where a flag marks the platform location. (F) Visual acuity in the optomotor test. Data for control (n=29 total, n=14 males, 15 females), *Adnp^Emx^*cHets (n=20 total, n=9 males, 11 females), *Adnp^Emx^* cKOs (n=20 total, n=9 males, n=11 females) and *Adnp^L822fs6/+^* gHets (n=14 total, n=7 males, 7 females) groups are plotted showing data for all mice (i), male mice only (ii), and female mice only (iii). Statistical analyses are via one-way ANOVA with the Tukey-Kramer post-hoc test for panel A. Statistical analyses are via two-way ANOVA with the Tukey-Kramer post-hoc test for panels B and C. \**p*□<□0.05; \*\**p*□<□0.01; *** *p*□<□0.001; \*\*\*\**p*□<□0.0001.

**Supplemental Figure 7. Additional analyses of the Morris Water Maze and visual tests.** (A) Average velocity during reversal probe day. (B) Frequency entering (right) and duration within (right) the quadrants of the Morris water maze during reversal probe day. (C) Frequency entering (right) and duration within (right) the platform target area during reversal probe day (Target FL), relative to the corresponding platform target areas in the other quadrants (Targets BL, BR and FR). (D) Schematic of platform reversal with example traces. (E) Latency to platform during Morris water maze visual test. (F) Averaged visual acuity measured via optomotor testing. Data for control (n=29 total, n=14 males, 15 females), *Adnp^Emx^*cHets (n=20 total, n=9 males, 11 females), *Adnp^Emx^* cKOs (n=20 total, n=9 males, n=11 females) and *Adnp^L822fs6/+^* gHets (n=14 total, n=7 males, 7 females) groups are plotted showing data for all mice (i), male mice only (ii), and female mice only (iii). Statistical analyses are via one-way ANOVA with the Tukey-Kramer post-hoc test for panel A. Statistical analyses are via two-way ANOVA with the Tukey-Kramer post-hoc test for panels B and C. \**p*□<□0.05; \*\**p*□<□0.01; *** *p*□<□0.001; \*\*\*\**p*□<□0.0001.

**Supplemental Figure 8. Conditional knockouts exhibit elevated freezing upon fear conditioning.** (A) Schematic of fear conditioning testing. During training (day 1) mice were placed in a novel cage environment. A tone was played three times while a mild electric shock was concurrently administered so the unconditioned context (cage environment) stimulus and cue (tone) stimulus were paired with the conditioned (shock) stimulus. On day 2, mice were re-introduced to the context (cage environment) of day 1 training. On day 3, mice were brought to a completely new context and the cue (tone) stimulus was played. (B) Percent of time inactive during training of unconditioned stimulus (shock) to conditioned context (cage) and cue (tone) stimuli. During “Tone-shock” phases, the shock was administered and tone cue was simultaneously played. (C) Percent of time inactive during the first three minutes versus the last three minutes of context testing, wherein mice were re-introduced to context stimulus (training environment) without shock. (D) Percent of time inactive prior to the cue, and post cue. Data for control (n=24 total, n=11 males, 13 females), *Adnp^Emx^* cHets (n=14 total, n=6 males, 8 females), *Adnp^Emx^* cKOs (n=14 total, n=5 males, n=9 females) and *Adnp^L822fs6/+^* gHets (n=11 total, n=6 males, 5 females) groups is plotted showing data for all mice (i), male mice only (ii), and female mice only (iii). Statistical analyses are via one-way ANOVA with the Tukey-Kramer post-hoc test for panel A. Statistical analyses are via two-way ANOVA with the Tukey-Kramer post-hoc test for panels B and C. \**p*□<□0.05; \*\**p*□<□0.01; *** *p*□<□0.001; \*\*\*\**p*□<□0.0001.

**Supplemental Figure 9. Detailed summary of ChaHP phenotypes integrating prior studies.** Phenotypic outcomes include viability, body and brain size measurements, exploratory behaviors, anxiety, repetitive behaviors, motor function, and learning. Stars indicate matched phenotypes with *Chd4^Nestin^* cKOs (Nitarska et al., 2016), as well as *Chd4^Emx^* cKOs and *Chd^Del/+^* gHets (Larrigan et al., 2023). *Adnp^Emx^* cHet and cKO neuroanatomical measures were previously reported (Clemot-Dupont et al., 2025). *Adnp^Del/+^*gHet behavioral data is preliminary and will be published elsewhere.

