## Supplementary figures and images for "An allelic series reveals the genetic requirement for Adnp in cortical neurogenesis and learning behavior"

### Fig, S4

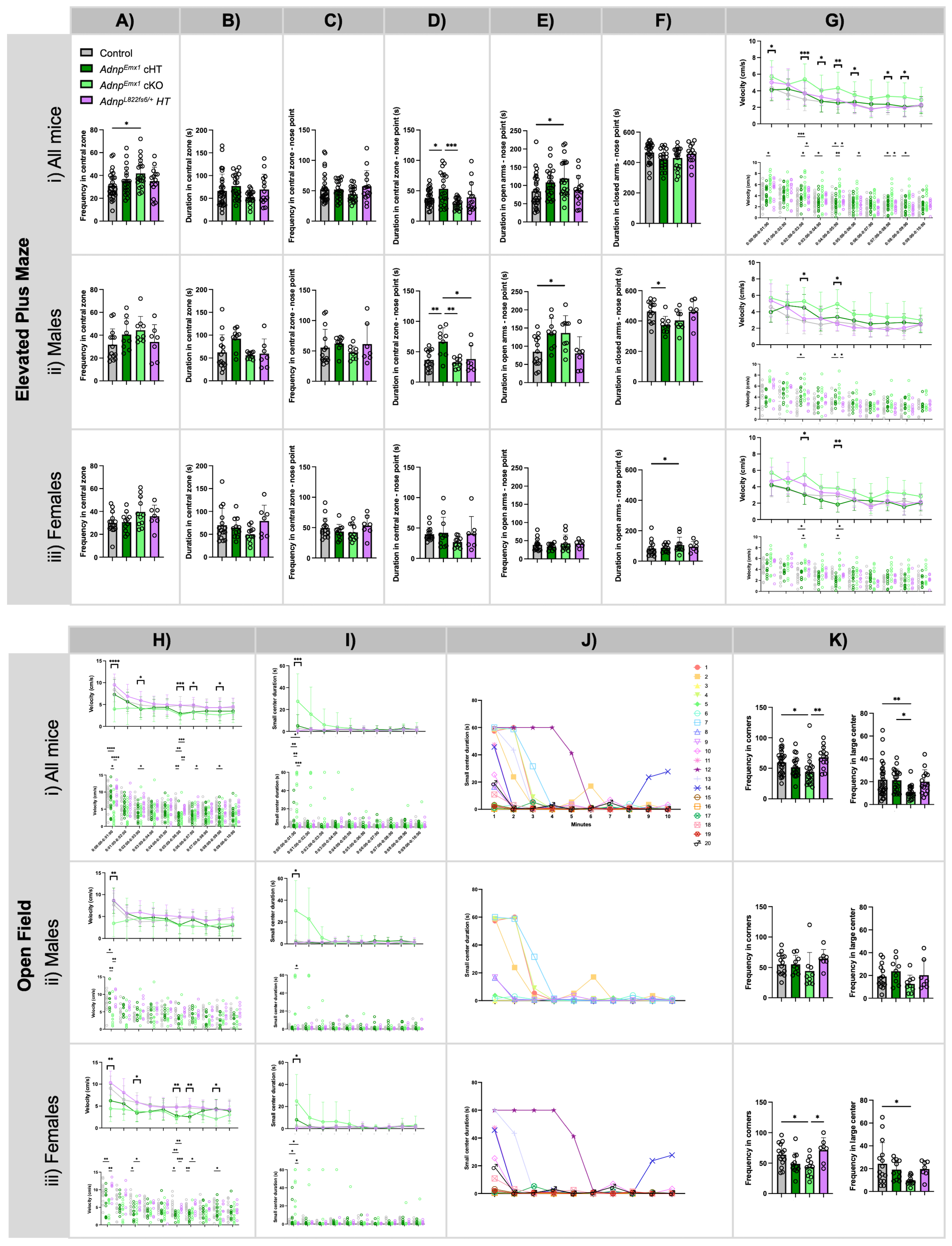

### Fig. S1

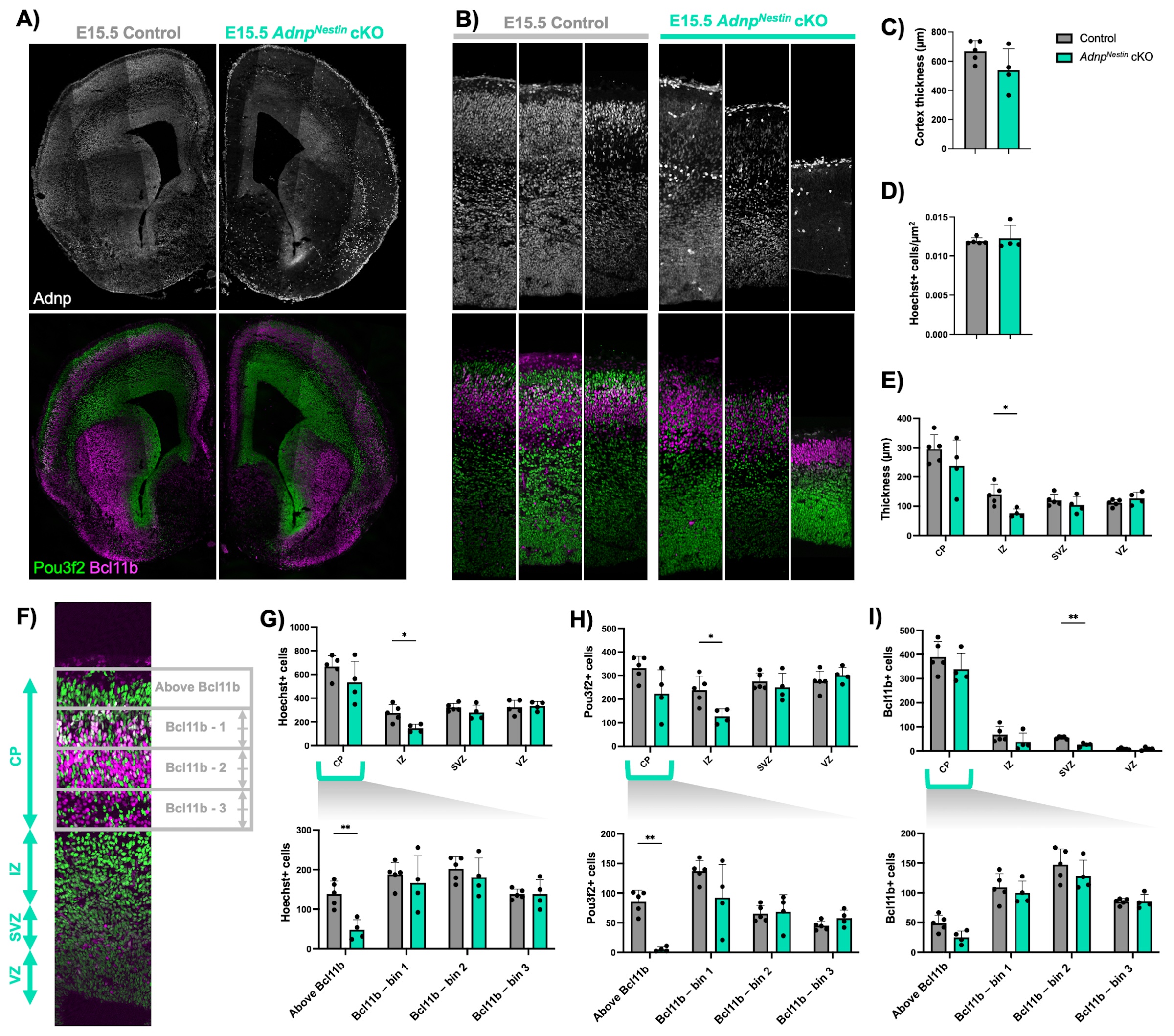

### Fig. S2

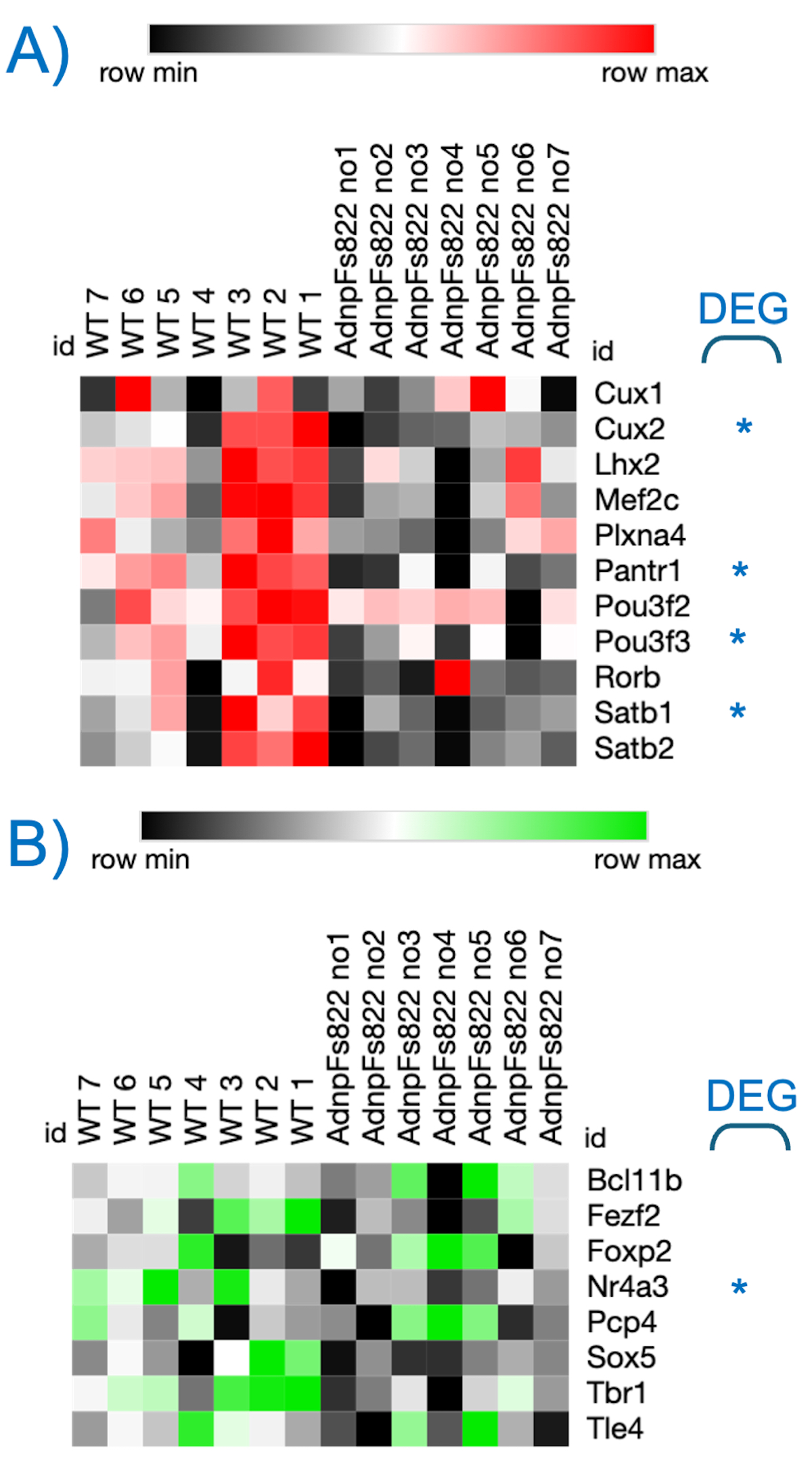

### Fig. S3

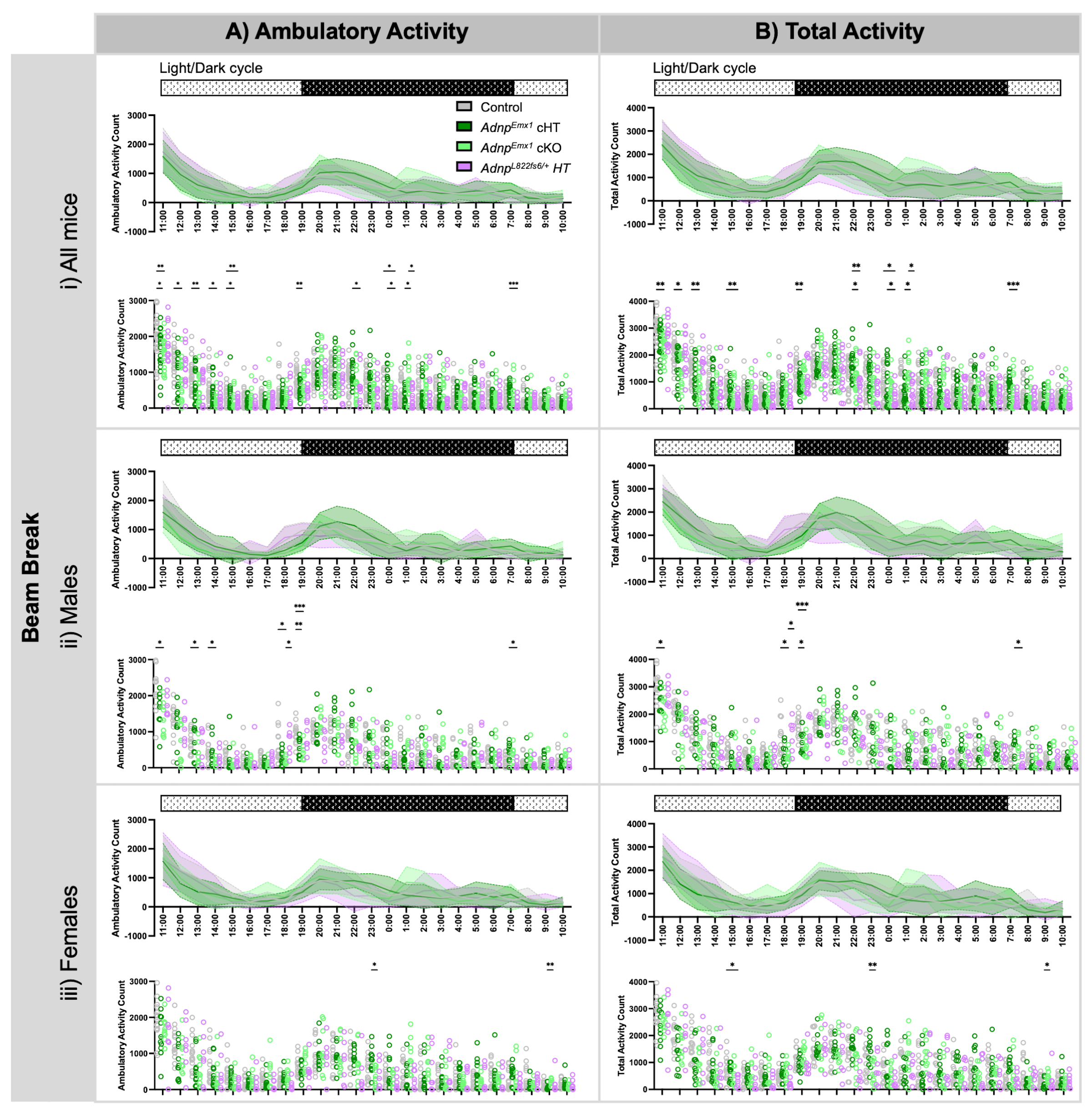

### Fig. S5

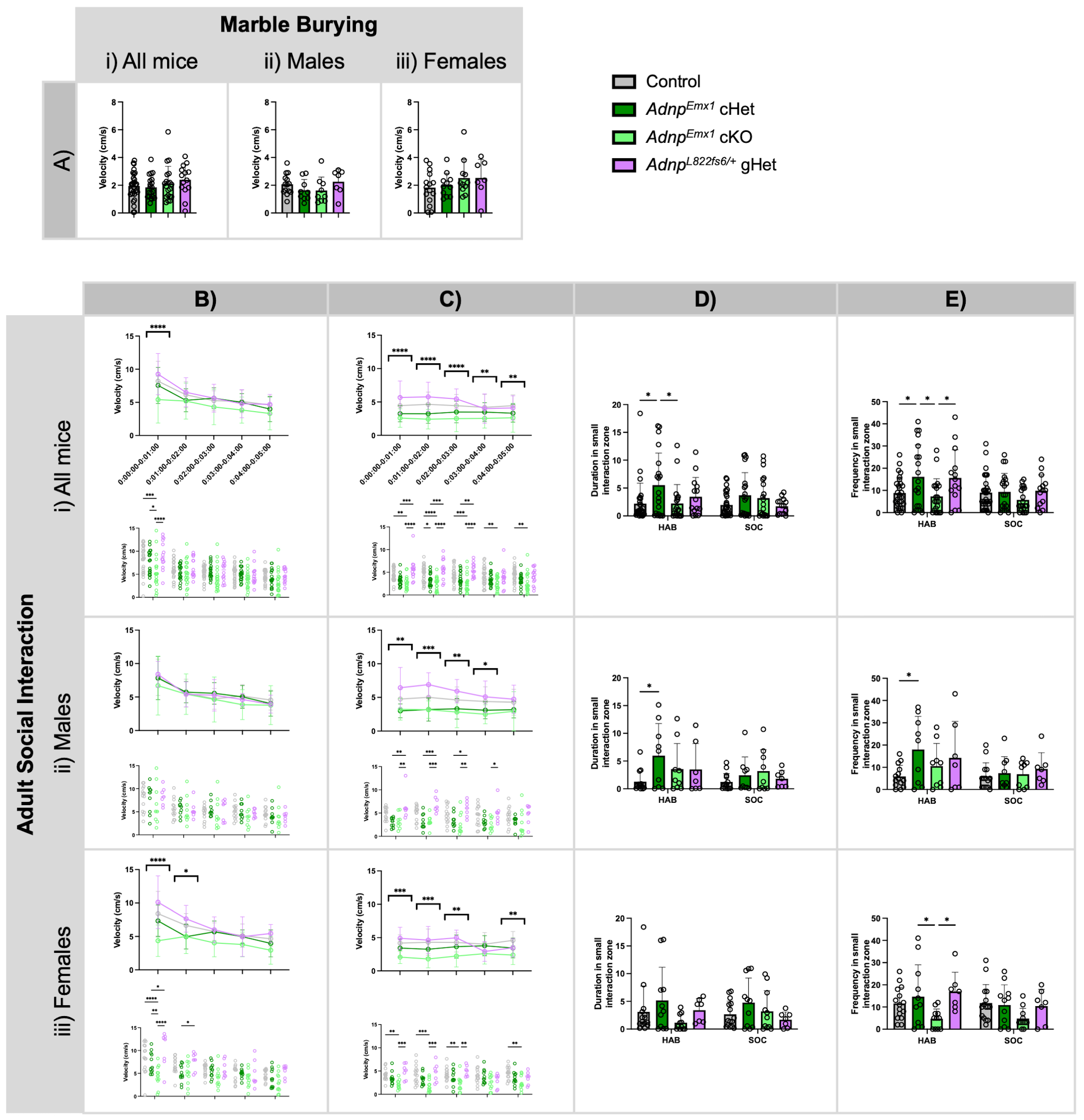

### Fig. S6

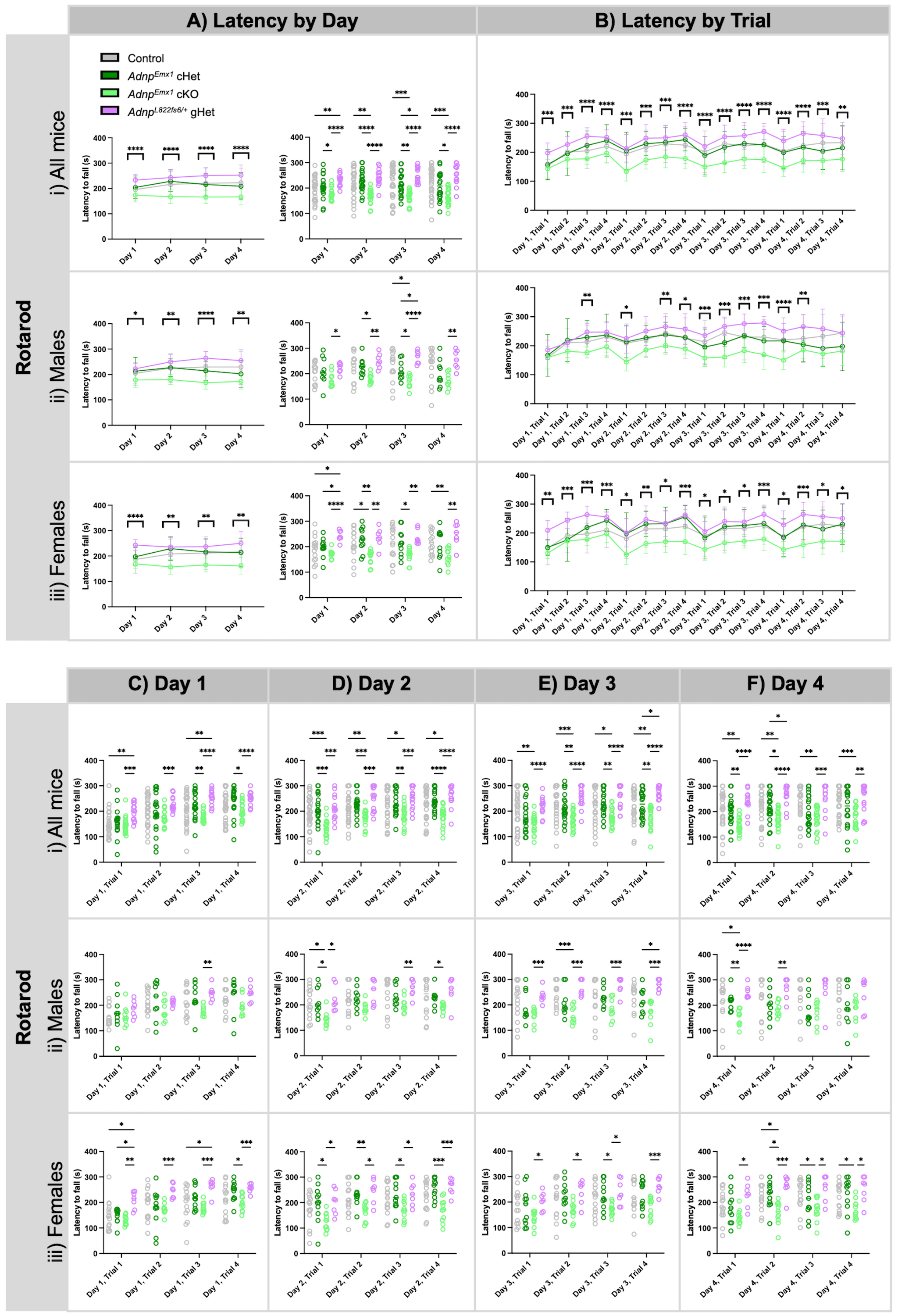

### Fig. S7

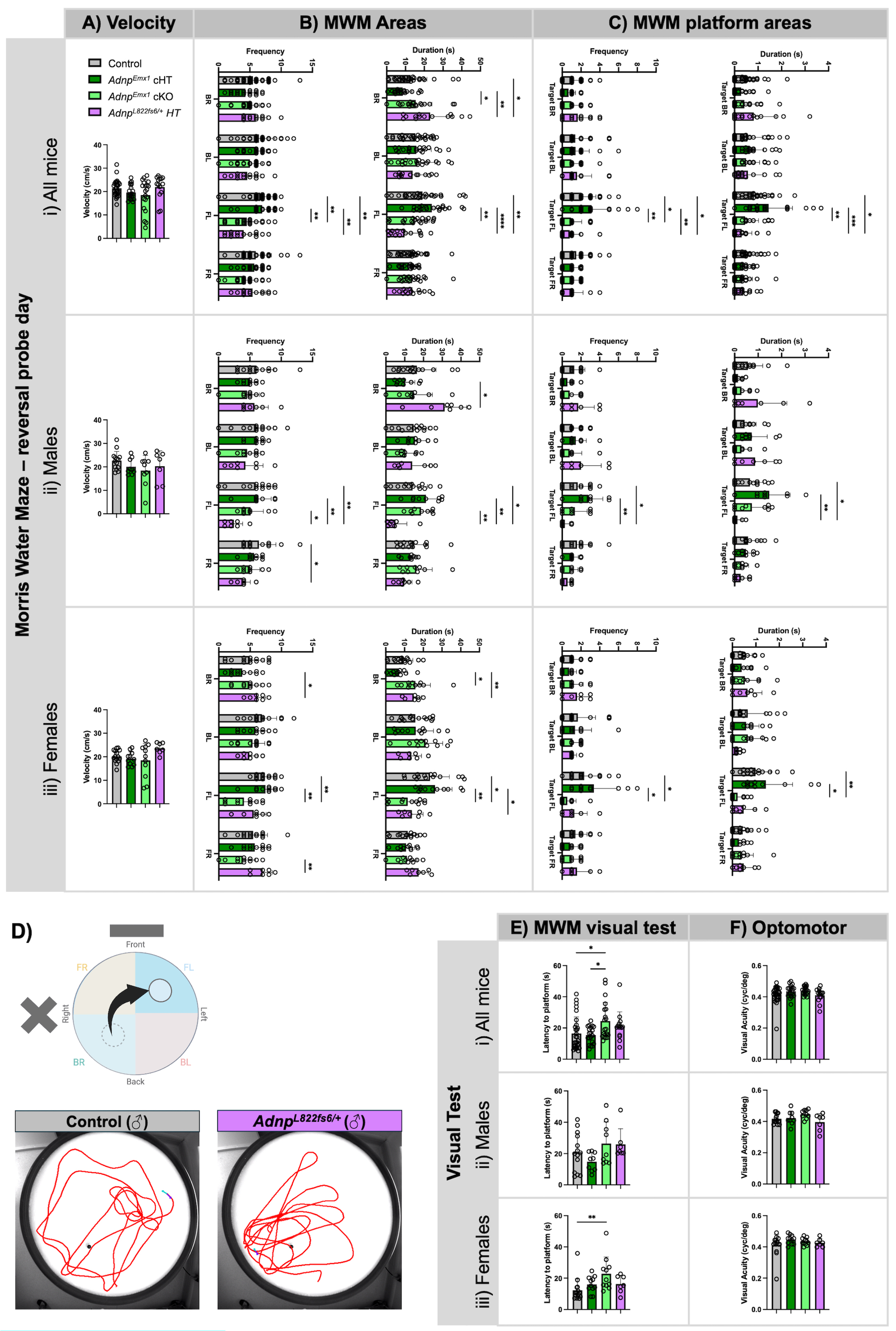

### Fig. S8

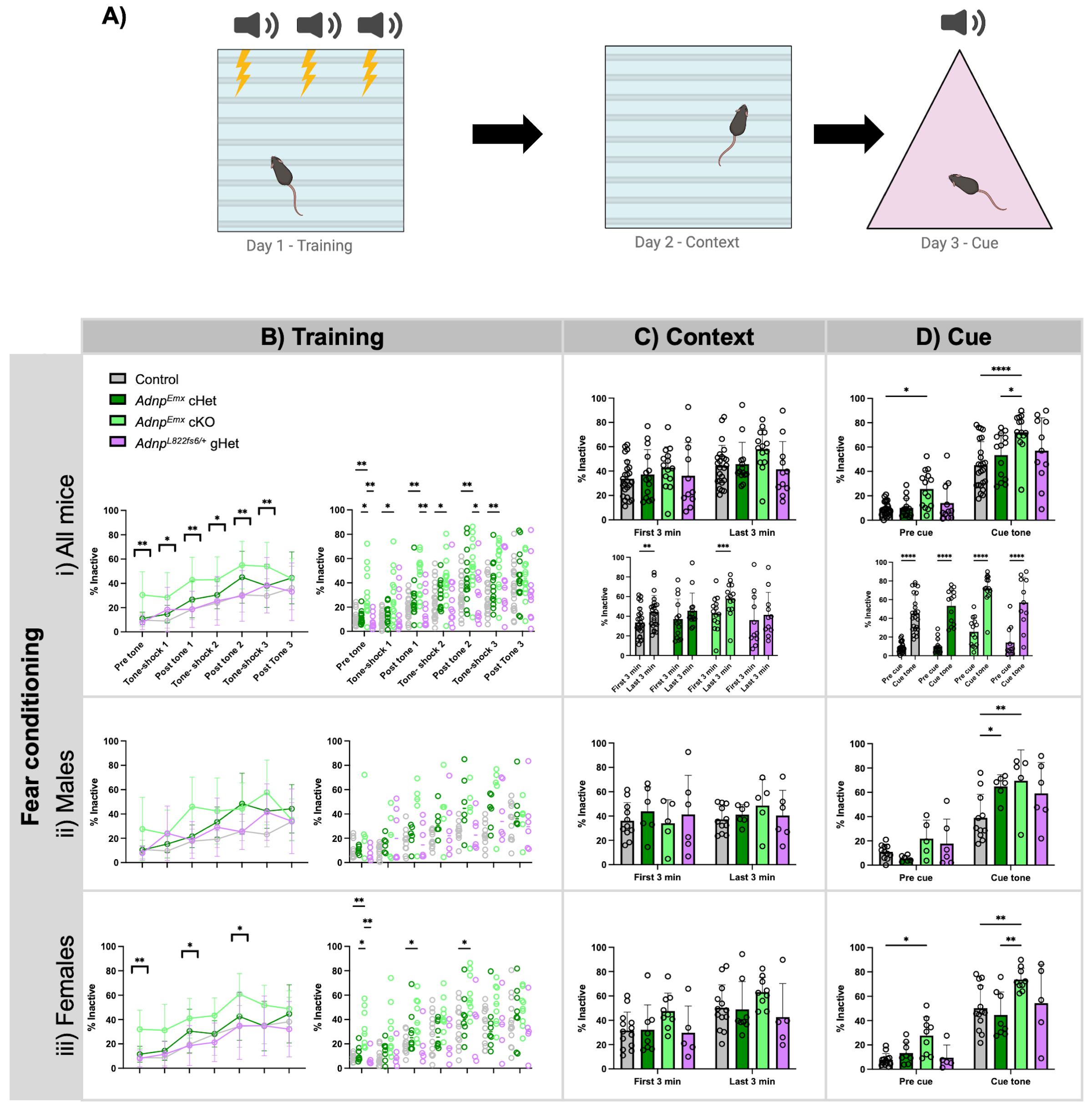

### Fig. S9

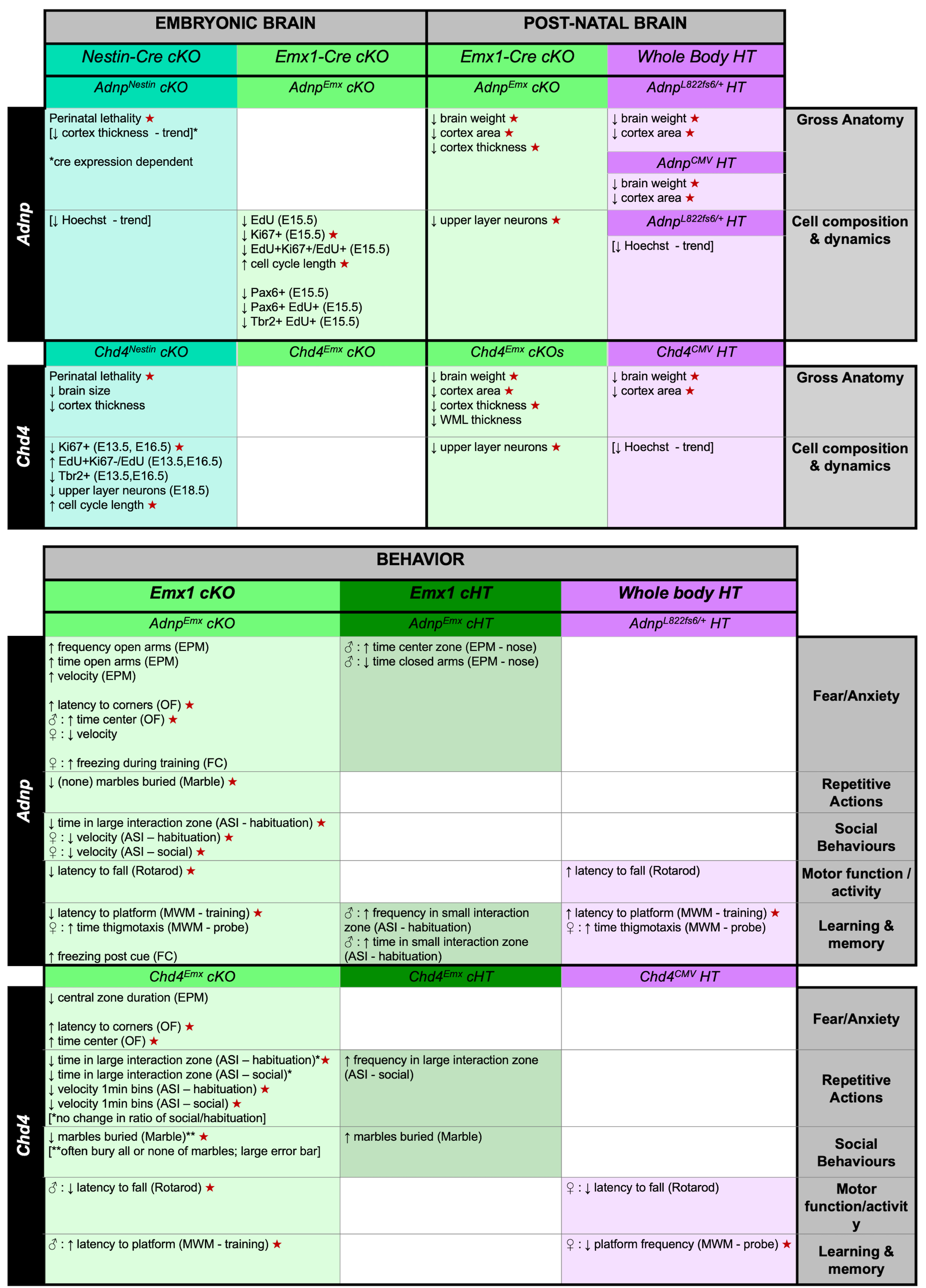
